# The influence of explicit local dynamics on range expansions driven by long-range dispersal

**DOI:** 10.1101/2022.09.15.508171

**Authors:** Nathan Villiger, Jayson Paulose

## Abstract

Range expansions are common in natural populations. They can take such forms as an invasive species spreading into a new habitat or a virus spreading from host to host during a pandemic. When the expanding species is capable of dispersing offspring over long distances, population growth is driven by rare but consequential long-range dispersal events that seed satellite colonies far from the densely occupied core of the population. These satellites accelerate growth by accessing unoccupied territory, and also act as reservoirs for maintaining neutral genetic variation present in the originating population, which would ordinarily be lost to drift. Prior theoretical studies of dispersal-driven expansions have shown that the sequential establishment of satellites causes initial genetic diversity to be either lost or maintained to a level determined by the breadth of the distribution of dispersal distances. If the tail of the distribution falls off faster than a critical threshold, diversity is steadily eroded over time; by contrast, broader distributions with a slower falloff allow some initial diversity to be maintained for arbitrarily long times. However, these studies used lattice-based models and assumed an instantaneous saturation of the local carrying capacity after the arrival of a founder. Real-world populations expand in continuous space with complex local dynamics, which potentially allow multiple pioneers to arrive and establish within the same local region. Here, we evaluate the impact of local dynamics on the population growth and the evolution of neutral diversity using a computational model of range expansions with long-range dispersal in continuous space, with explicit local dynamics that can be controlled by altering the mix of local and long-range dispersal events. We found that many qualitative features of population growth and neutral genetic diversity observed in lattice-based models are preserved under more complex local dynamics, but quantitative aspects such as the rate of population growth, the level of maintained diversity, and the rate of decay of diversity all depend strongly on the local dynamics. Besides identifying situations in which modeling the explicit local population dynamics becomes necessary to understand the population structure of jump-driven range expansions, our results show that local dynamics affects different features of the population in distinct ways, and can be more or less consequential depending on the degree and form of long-range dispersal as well as the scale at which the population structure is measured.

## I. INTRODUCTION

Range expansion—the act of a population expanding into new territory—is common in biological populations. Range expansions occur naturally and randomly all the time, often as the result of a species’ natural movement, such as by animals moving into new territory or maple helicopters carrying seeds away from their parent tree. Researchers have documented range expansions in a wide variety of organisms, such as plants [1], birds [2], sea creatures [3, 4], and terrestrial animals [5, 6], even humans [7]. Range expansions are increasingly forced by global warming as the changing climate makes traditional habitats inhospitable, while potentially opening up new hospitable regions [8].

Range expansions leave distinctive signatures in the patterns of genetic diversity of a population that can mimic the effects of natural selection [9]. Individuals at the frontier of an expanding population make a large contribution to the subsequent expansion wave, even if their frontier position was solely due to chance; as a result, genetic variants they carry can acquire high frequencies in the population in a phenomenon termed gene surfing [10, 11]. Independent surfing events in separate sections of the expansion front cause the population to segregate into genetically distinct sectors, promoting an illusion of local adaptation from purely neutral mutations [12–15]. Modeling the combined effect of spatial structure and stochasticity on neutral genetic diversity is key to understanding the biological origins of established genetic patterns, and to the successful prediction of future genetic diversity in pandemics and ecological expansions.

The influence of random chance on genetic diversity during range expansions can be amplified by long-range dispersal [16]. Many species have evolved ingenious ways of dispersing offspring over long distances with help from natural forces and from other organisms [17]. Plants rely on the dispersal of seeds and pollen by wind, waves, and animals [18]. Glacier ice worms can travel hundreds of miles, likely carried by migratory birds [19]. Modern pandemics are driven by microorganisms hitchhiking on air travelers to find new uninfected populations [20]. Even if long-range dispersal events are rare, they have an out-sized influence on the expansion because they enable pioneers to seed satellite colonies in uninhabited areas. If a pioneer happens to land in a place with abundant resources and little to no competition, its descendants may flourish. The pioneer’s genes will then propagate and any genetic variants they carry will reach high frequencies in the vicinity of the satellite [21–25] even in the absence of a selective advantage; random chance alone has caused the pioneer’s genes to become prominent by means of a founder effect, leading to a *suppression* of local diversity within satellites. However, long-range dispersal also *favors* neutral diversity at larger scales, by ensuring that individuals well within the expanding population have a chance of contributing to growth. The evolution of overall diversity during the range expansion is governed by the trade-off between the two effects, and can depend sensitively on the degree of long-range dispersal experienced by the population [26, 27].

Modeling the general characteristics of range expansions requires two minimal ingredients: a probability distribution of dispersal distances *J* (*r*), also called the *jump kernel*, from which dispersal events are randomly drawn; and a method of local density regulation to model the existence of a finite carrying capacity. When long-range jumps are present, the tail of the jump kernel, i.e. its behavior at long distances, critically influences the fate of the population at long times. Fundamental differences from short-range dispersal are observed when the jump kernel is “fat-tailed”; i.e. it decays slower than exponentially with increasing distance. Fat-tailed jump kernels lead to expansions that accelerate as they progress, unlike the constant-speed expansions that occur when dispersal is exclusively short-range [28].

A commonly used fat-tailed kernel is the power-law jump kernel *J* (*r*) ∼ 1*/r*^*μ*+1^. Besides providing a good description of the dispersal behavior of many species [29], power-law kernels are a useful tool for analyzing and classifying the breadth of potential population outcomes due to long-range dispersal [16]. The exponent *μ* is a key factor governing the long-time characteristics of the growth and the dispersal patterns, whereas other details of *J* (*r*) such as its short-distance functional behavior are less relevant [16]. A broad range of expansion behaviors is encompassed by varying the kernel exponent (limited to *μ* > 0 to ensure a normalizable distance distribution). At high *μ*, the jump kernel decays quickly with increasing distance, and a colony expands at a constant rate as if there were exclusively short-range dispersal. As *μ* → 0, spatial structure becomes irrelevant and a colony grows as if it were in a well-mixed liquid environment. The intermediate range of kernel values connects these two extremes in a tunable manner.

Recent work has catalogued the distinctive features of population growth dynamics [16] and spatial genomic patterns [27] that can be achieved upon varying the kernel exponent in range expansions driven by power-law growth kernels (a detailed summary is provided in Section II). These studies have identified a critical value of the kernel exponent *μ* below which the population grows nearly as fast as a well-mixed population, and a significant fraction of the neutral variation in the originating population is preserved for arbitrary long times due to serial reintroduction of variants from the core of the expanding population. For kernel exponents close to but above the critical threshold, population growth is slowed down dramatically and neutral diversity is steadily eroded. However, at even higher values of *μ*, the behavior approaches that of short-ranged jump kernels, where the population advances as a front moving outward at constant speed. In this situation, a small fraction of the diversity in the originating population persists due to the formation of sectors [12, 30].

Less well understood is the influence of the second key feature of spatial population models: the density regulation mechanism. Modeling growing populations in a spatial continuum presents challenges to both the forward-in-time [31] and backward-in-time [32, 33] approaches, due to the necessity of systematically imposing a local region of influence within which each individual can impact the growth of its neighbors. Local density regulation is commonly implemented by dividing up space into a regular grid of well-mixed subpopulations called demes, each of which has a fixed carrying capacity. Migration events, drawn from the jump kernel, transport individuals across demes. Deme-based models and their variants are widely used in population genetics [34], including for the study of range expansions [9, 30]. However, models that rely on a lattice of demes have their limitations. By design, they do not capture spatial structure and stochasticity at scales smaller than the effective deme size. Imposing an artificial grid of demes also introduces artifacts to the population structure, which can in some instances get *worse* upon increasing the grid resolution to better approximate a continuum [35].

Additionally, using deme-based models forces researchers to make decisions about the specifics of deme saturation and population management. The following selection of recent work exemplifies various possible strategies. Some may choose to have demes that instantaneously change from being empty to full upon the arrival of the first migrant [16, 27], while others may let the deme population grow logistically at a predetermined rate [36, 37] or let the growth be determined by random migration events that bring in individuals from other demes [38]. Death can occur in various ways, such as by attempting to disperse into an already full deme [16, 27] or by being randomly resampled out of an overfull deme’s population [38]. If the density regulation unit is the deme population as a whole rather than the individuals in the deme, death may not explicitly occur to any individuals, but the deme population size changes from one time step to the next [36, 37]. Since most computational studies involving long-range dispersal, including the quoted prior results [16, 21–25, 27, 36, 37, 39], have relied on deme-based approximations, the applicability of their conclusions to continuum-space population growth remains an open question.

The aforementioned results on the population dynamics and neutral evolution of range expansions driven by power-law kernels [16, 27] were derived using a lattice of demes with an additional simplifying assumption: upon arrival at an empty deme, the pioneer immediately saturates the deme, excluding any other migrants from establishing themselves. Not only does this assumption exclude any effects of local dynamics on population growth, it also enforces a local *founder takes all* effect where only one migrant is allowed to contribute to the genetic makeup of a density regulation region. Instant local saturation is justified when long-range jumps are rare and most offspring land within a short distance of their parents; then, the local logistic growth within a deme occurs extremely fast compared to the typical time to arrival of another migrant from a different deme, and can be treated as instantaneous. However, the instant saturation and founder-takes-all assumptions can be invalid when the time scales of local and long-range dispersal are comparable, in which case a local region might receive and send out several migrants while it is being saturated. The influence of the breakdown of fast local saturation on the population dynamics and the spatial genomic structures left behind by long-range dispersal is unknown.

In this work, we address these gaps in our knowledge of range expansions driven by long-range dispersal by performing and analyzing continuum space, individual-based simulations of range expansions driven by power-law kernels. Our simulations were implemented in the population genetics program SLiM [40], and do not use a grid of demes or assume instant saturation of the local carrying capacity by the first arrival. Instead, individuals occupy positions in continuum space and their survival depends on the number of other individuals present within a defined region of influence at the time of their birth. When possible, we compared the outputs to the predictions from models based on lattices of demes of the population growth rate [16] and the evolution of neutral genetic diversity [27]—we term these prior models “lattice-based” predictions. We found that our results often agreed with the lattice-based predictions, giving conditional support to prior results based on models that only focus on the founders. However, when individuals can share resources with many others, we found that focusing exclusively on the founders misses important dynamics between coexisting or competing alleles. In those cases, it becomes necessary to also consider individuals who arrive after the pioneer. We identify parameter regimes where using the lattice-based models is justified, and show that they depend on the specific kernel exponent.

## II. BACKGROUND

We first summarize prior results [16, 27] on range expansion dynamics for populations experiencing long-range dispersal with fat-tailed kernels, which were obtained using lattice models conforming to the founder-takes-all assumption at the deme level. Ref. 16 used a lattice model to quantify how a colony expands into unoccupied space when offspring are dispersed according to a power law jump kernel that decays according to *J* (*r*) ∼ 1*/r*^*μ*+1^. The authors showed that the power-law tail captures the qualitative features of the long-term population growth, and that the short-range behavior of the jump kernel has a negligible impact on the long term population growth. The model of Ref. 16 (hereafter “the lattice model”) divides *d*-dimensional space into a lattice of habitats or “demes”. Occupied demes generate offspring according to a Poisson process; offspring attempt to migrate to a new deme randomly chosen by drawing a dispersal distance from *J* (*r*) and a random direction relative to the originating deme. Instant local saturation is assumed and is enforced in the model by allowing only two states to each deme: occupied, or empty. A migration attempt to an empty deme is successful, and immediately turns the state of that deme to occupied. A migration attempt to an occupied deme is unsuccessful, and the offspring dies. These assumptions guarantee a founder-takes-all effect at the local level. Henceforth, when we refer to the lattice model, it is implied that instant local saturation and local founder-takes-all are enforced. Much of our current understanding of jump-driven range expansions derives from the lattice model, as summarized below.

### A. Population growth and time-doubling hierarchy

Analysis of the lattice model [16] showed that at all times *t*, a core region of the colony can be identified that is centered at the originating population of the range expansion and within which most demes are occupied. The size of this core region is proportional to the total population size *M* (*t*) of occupied demes in the expansion. The long-time asymptotic behavior of the radius of the core region ℓ(*t*) ∝ [*M* (*t*)]^1*/d*^ depends on the “heaviness” of the tail of the jump kernel, which is set by the kernel exponent *μ*. There are two distinct growth possibilities, separated by the value *μ* = *d* + 1: the colony expands at a constant rate for *μ* > *d* + 1 and it expands faster than linearly when *μ* < *d* + 1 [16]. The faster-than-linear growth regime is driven by long jumps whose characteristic size continues to increase as the core expands: this “jump-driven” growth regime, in which pioneers have a large impact on growth, will be the focus of this paper. Within the jump-driven regime, a second special value *μ* = *d* separates two distinct asymptotic behaviors of the core growth at long times (*t* → ∞): when *d* < *μ* < *d* + 1, the core grows asymptotically as a power law which is faster-than-linear in time 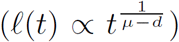, in contrast to stretched-exponential growth when *μ* < *d* (ℓ(*t*) ∝ exp(*B*_*μ*_*t*^*η*^), where *B*_*μ*_ and *η* themselves depend on *μ* and *d*).

A key result of Ref. 16 was that expansions in the jump-driven regime are governed by a hierarchical time-doubling structure, as depicted in Fig. 1. The core of the colony expands by “absorbing” satellite colonies that were seeded at an earlier time by a rare but consequential long jump. A typical satellite being absorbed into the core at time *t* was seeded approximately at time *t/*2 by an offspring who dispersed roughly a distance ℓ(*t*) from its parent in the core of the colony. Mathematically, this self-consistency condition can be expressed as

**FIG. 1.**
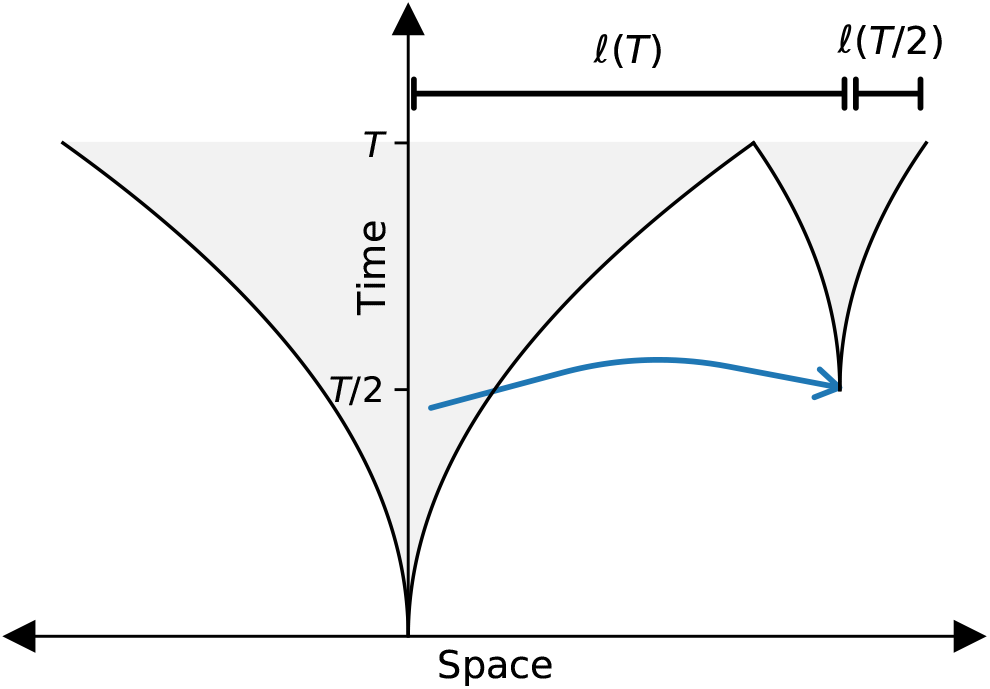
Schematic diagram of the time doubling hierarchy discovered by Ref. 16. Shaded parts of the plot represent regions of space that are occupied at a given time. The core of the colony (central funnel) grows by absorbing satellites that were seeded at an earlier time by long-range dispersal. A typical satellite being absorbed into the core at time *T* (smaller funnel at right) was seeded at time of order *T/*2 by an offspring who dispersed a distance of roughly ℓ(*T*) from its parent in the core of the colony; it has grown to a size of order ℓ(*T/*2) when it merges with, and becomes part of, the core.

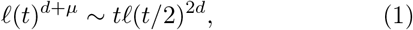

where the tilde signifies agreement of the leading functional dependence of either side of the relation on the time variable, without including time-independent prefactors or terms whose fractional contributions vanish at long times. The time-doubling hierarchy and Eq. (1) form the basis for deriving the asymptotic functional forms of ℓ(*t*) summarized above; unlike those asymptotic forms that are valid only at very long times *t* → ∞, the self-consistency condition holds as long as the population is large enough that an appreciable number of long-range jumps have occurred [16]. Equation (1) forms a basis for more accurate functional forms of the outbreak growth dynamics [16], and also leads to quantitative insights into the evolution of genetic diversity when multiple variants are present in a population experiencing long-range dispersal [27, 39]. Note that the time-doubling hierarchy only relies on the assumption of instant local saturation, and does not require that space be discretized into a regular lattice of demes.

### B. Persistence of initial neutral variation

A striking consequence of range expansions is that the combination of stochasticity and spatial structure can leave behind patterns of neutral genetic variation that are typically associated with selection, such as sweep-like enrichment of individual alleles [10], diversity gradients [11], and segregation of variants into distinct regions [12]. Simplified models of neutral evolution in spatially structured populations enable us to understand such patterns and to distinguish them from the outcomes of selective events. One aspect of neutral variation that is closely tied to the mode of dispersal is the persistence of initial genetic diversity in the originating population during its expansion into new territory [21, 26]. When dispersal is exclusively short-ranged, only individuals near the edge of the range expansion contribute to future variation; in the absence of new mutations, much of the initial diversity can be lost over time due to successive founder events at the edge. Long-range dispersal enables regions far within the population to contribute to the expansion, which maintains their alleles in the growing population and favors diversity. However, founder effects are not eliminated: each long-range jump seeds a satellite outbreak in which all offspring share the allelic identity of the seeding pioneer, acting as a genetic bottleneck which eliminates diversity locally in the absence of mutations. The fate of the initial neutral variation as the expansion progresses is determined by the balance between these contrasting effects.

The evolution of initial neutral diversity in jump-driven range expansions was analyzed in Ref. 27, using a lattice model in which neutral variation was introduced in the starting population, and no new mutations appeared during the expansions. The existence of the time-doubling hierarchy, Eq. (1), was used to identify an effective population of homogeneous satellites whose evolution captures the balance between diversification and coarsening for a given jump kernel exponent. As with the behavior of the core radius growth, the amount of initial diversity preserved after a range expansion was shown to suffer different fates depending on the value of the kernel exponent relative to the spatial dimension. When *μ* < *d*, the diversifying influence of long jumps dominates; note the large number of satellites well separated from the core in Fig. 2**g–i**. The seeding of many satellites by long-range dispersal events from the core enables the population to preserve a finite amount of its initial heterozygosity at long times. By contrast, when *d* < *μ* < *d* + 1, the local coarsening of diversity due to bottlenecks becomes more significant; note the small number of large monoclonal satellites in Fig. 2**d–f**. The heterozygosity decays inexorably towards zero as the range expansion progresses, albeit at a slow rate. As *μ* approaches *d*, the heterozygosity approaches a finite value but the convergence to this value becomes extremely slow and cannot be observed over practical simulation times. Notably, for *μ* > *d* + 1, some diversity is also preserved at long times due to the formation of sectors in outward range expansions, as shown in Fig. 2**a–c** [12, 30]. Jump kernels of intermediate breadth (*d* < *μ* < *d* + 1) therefore support lower neutral diversity than broader (*μ* ≤ *d*) and narrower (*μ* ≥ *d* + 1) kernels.

**FIG. 2.**
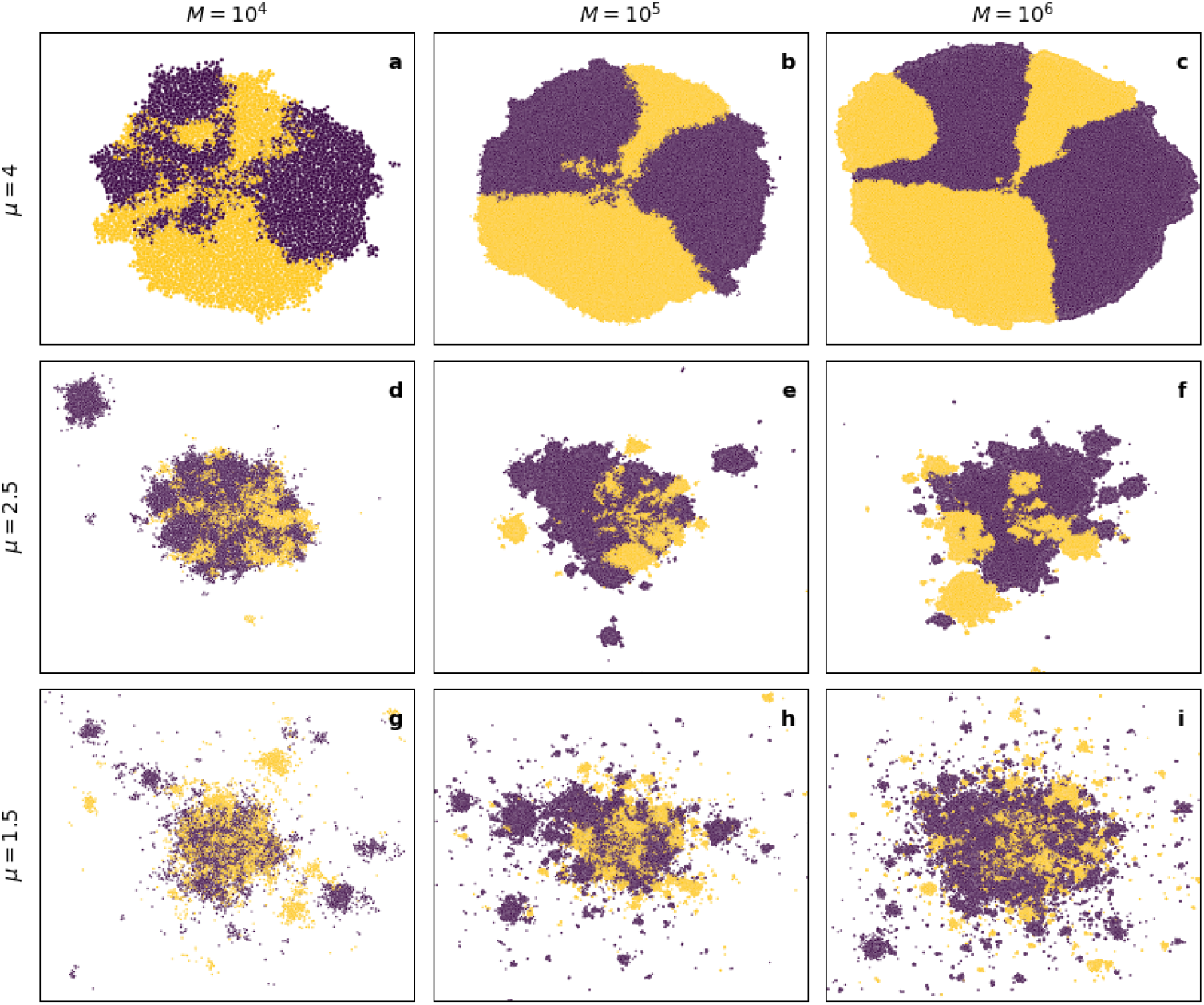
Snapshots of simulated range expansions at different population sizes *M*. These simulations began with 100 individuals equally split between two neutral alleles, labeled as either purple or yellow in these plots. **a-c.)** Diversity is preserved by the formation of monoallelic sectors for *μ* > *d* + 1. **d-f.)** The small number of satellite outbreaks act as bottlenecks, eroding diversity for *d* < *μ* < *d* + 1. **g-i.)** Long-range jumps transport alleles from the core to the exterior of the colony, preserving diversity for *μ* < *d*. Additional parameters are *K* = 10 and *p* = 0.

In summary, Ref. 27 established that long-range dispersal can preserve some of the genetic diversity from the originating population at long times, but only for jump kernels broader than a dimension-dependent threshold. Narrower kernels cause diversity to erode over the course of the expansion due to successive founder events, which can erase even the limited heterozygosity preserved due to the formation of sectors in range expansions with exclusively short-ranged dispersal. However, these features were observed in lattice models which assumed instant local dynamics and founder-takes-all at the deme level; the influence of slow local saturation on the evolution of heterozygosity could not be gauged. In this study, we aim to establish whether insights derived from lattice models of range expansions still apply in a continuous-space model for which the lattice model assumptions can be violated to a controllable degree, and to quantify the effect of explicit local dynamics on neutral genetic variation as the expansion progresses. We next introduce our simulation model which we use to investigate these questions.

## III. METHODS

In order to study jump-driven range expansions which rely on neither a lattice nor the assumption of instant local dynamics, we used the evolutionary simulation software SLiM [40] to simulate range expansions on a 2D continuous landscape without restricting ourselves to a lattice of demes. Individuals produce offspring at a constant rate, and offspring attempt to establish themselves by dispersing in a random direction with dispersal distances drawn from a jump kernel incorporating short-ranged and long-ranged dispersal which we define below (Eq. (3)). To focus on the effects of the spreading process and enable direct comparison with previous work (see Section II), our model includes two simplifying assumptions. First, each individual has an allelic identity which is passed on exactly to offspring with no possibility of new mutations; this enables us to evaluate the persistence of initial neutral variation purely due to dispersal and spatial structure during spreading. Second, once offspring are successfully established, they do not move, die, or renew themselves. This assumption allows us to hone in on the dynamics of establishment and expansion, without confounding effects or computational expense from reshuffling and replenishment of regions that have already been saturated. Immortality and immobility post-establishment provide a reasonable approximation for trees that produce massive numbers of seeds over scores of growing seasons, or perennial plants that replenish themselves in place once established. Even in populations for which these assumptions do not hold, the patterns left behind by the initial expansion can still be representative of long-time trends despite the subsequent gene flow due to replenishment and reshuffling of individuals [14, 15].

In the absence of demes with a fixed carrying capacity, a different mechanism to regulate population growth is needed. We assume that the environment has uniformly-distributed resources which can support a uniform carrying capacity per unit area, quantified by a maximum population density *ρ*. We introduce an interaction distance *r*_b_ which demarcates a disc-shaped region within which an individual competes with others for resources (Fig. 3**a**). The population density and interaction distance can be combined to define a local carrying capacity *K* via

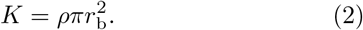

When an individual is born, it undertakes a random dispersal event and counts the number of individuals within the interaction region surrounding its new location. If there are at least *K* other individuals in the interaction region, the duplication event is unsuccessful and the new individual dies. If there are fewer than *K* other individuals, the new individual establishes successfully in its new location and survives for the remainder of the simulation.

**FIG. 3.**
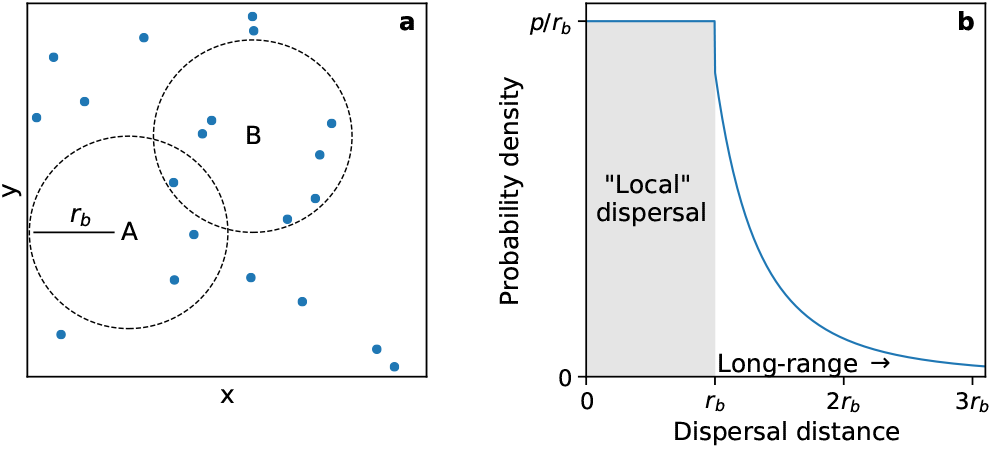
An outline of the simulation procedure. **a.)** A snap-shot of a population during a range expansion. The dots represent individuals in the population. Suppose the local carrying capacity is 5. An individual born at position *A* would only count three others in its local region (dashed circle centered at *A*), so it would survive. An individual born at position *B* would count seven others within its local region. That is too many for the individual to successfully compete against, so it would die. **b.)** An example jump kernel. There is a probability *p* of dispersing within the “local” (shaded) region, that is, within distance *r*_b_. The jump kernel decays according to the power law *J* (*r*) ∼ 1*/r*^*μ*+1^ beyond the local region.

The local interaction region in our continuous-space simulation resembles the geographic subdivision unit (the deme) used in lattice-based models. The concept of instantaneous local saturation, or a local founder-takes-all effect, would therefore correspond to an individual quickly filling its interaction region with its offspring before it (or its descendants) attempted any long-range dispersal events. In order to smoothly depart from the assumptions of the lattice model, it would be useful to control the fraction of dispersal events which are “local”, i.e. within the interaction region, as opposed to long-range. To do so, we used a two-part jump kernel that allows us to explicitly specify the probabilities of local versus long-range dispersal, as sketched in Fig. 3**b**. In full, the jump kernel is as follows:

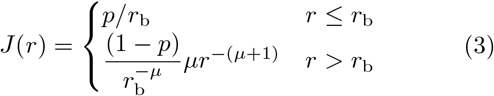

where *p* is the probability of dispersing within the local region. The short-range part of the jump kernel is chosen to be featureless, with the only notable property being that the integrated probability 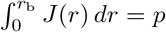. The long-range part of the jump kernel matches the power-law kernel used in the prior works discussed [16, 27, 39] and the prefactor ensures the normalization 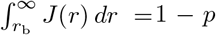. Jump distances are randomly drawn from this distribution using inverse transform sampling (detailed procedure in SI Section VI A).

A few comments about our choice of jump kernel, Eq. (3), are in order. Our aim is not to exactly reproduce a biologically measured jump distribution at all lengths, but rather to capture the two main features of interest in a simplified kernel—a tunable balance between short- and long-range dispersal determined by the parameter *p*, and a fat-tailed kernel with a specified power-law falloff controlled by the exponent *μ*. For simplicity, we chose the short-range part of the jump kernel to be constant with distance *r*; other forms are expected to lead to similar results provided the integrated probability of jump lengths between 0 and *r*_b_ evaluates to *p*. The chosen form also implicitly assumes that the same length scale *r*_b_ governs the interaction distance for the density regulation and the dispersal behavior. We could have built a model with an additional length parameter dictating the spatial features of the dispersal kernel, but at the cost of added complexity and a larger parameter space. Our simplified choice allows us to dial in a specific balance between local and long-range dispersal by adjusting the parameter *p* alone, which enables direct comparisons of different simulations where the kernel exponent, local carrying capacity, and size of the density regulation region are kept unchanged. Since the exact shape of the jump kernel at short distances is not biologically realistic (for instance, it has a discontinuity at *r* = *r*_b_), we do not use our model to draw any conclusions about the spatial distribution of individuals on scales smaller than the interaction distance.

We now specify appropriate units for length and time in our simulations. Since the individuals and the environment are both featureless, and the same length scale *r*_b_ governs both the density regulation and the dispersal, the interaction distance is the natural length unit in our model. In our simulations, we set *r*_b_ to one, so that all distances reported from simulations are in units of *r*_b_. Time units are chosen such that each individual generates offspring via a Poisson process with a duplication rate of one; i.e. time is reported in units of the average generation time for an individual. Note that not all offspring survive, because of the density regulation mechanism.

Once the length and time units have been fixed, the consequential parameters are the kernel exponent *μ*, the probability of local dispersal *p*, and the local carrying capacity *K* (which determines the local density *ρ* via Eq. (2)). Simulations begin with 10*K* individuals whose *x* and *y* positions are random draws from a Gaussian distribution with mean zero and standard deviation 2*r*_b_. Everyone in the population gets a chance to produce offspring every time step, which disperse according to the jump kernel with relevant *p* and *μ* and then either survive or don’t depending on the population density where they happen to land. Simulations end once the population size exceeds a predetermined threshold, usually four orders of magnitude larger than the initial population size. See SI Section VI A for more details on the simulation procedure.

We next identify characteristic time scales in the problem which will enable us to choose parameters which violate the instant local dynamics and local founder-takes-all assumptions. (For an expanded discussion with potential improvements, see SI Section VI B). First let us consider the characteristic saturation time scale for a single interaction region (which takes the place of a deme in our model). While the full saturation dynamics is complicated because of the influence of offspring from nearby interaction regions, we can make a simplified estimate of the saturation time by considering only the descendants of the pioneer individual which undergo local dispersal. Assuming that all these descendants land in the same interaction region, we have an effective division rate of *p* (in our units) for the local population. In this simplified model, the interaction region fills up according to a logistic function with growth rate *p*, for which the saturation dynamics are set by the characteristic time scale *τ*_s_ ≡ 1*/p*. (The actual saturation time for a deme with a discrete population has an additional logarithmic dependence on the carrying capacity, see SI Section VI C; we ignore this weaker dependence compared to the dominant 1*/p* dependence in the present discussion of characteristic time scales.) The saturation time scale must be compared to the typical time for the interaction region to send out long-range jumps. The highest possible rate occurs when the region has saturated to population *K* and sends out long-range jumps at a rate *K*(1 − *p*). Therefore, we identify *τ*_j_ ≡ 1*/*(*K*(1 − *p*)) as the characteristic time scale separating long-range jumps out of an interaction region. Note that the local saturation time scale is independent of *K*, whereas the rate of long-range jumps out of an interaction region does depend on *K*.

The instantaneous local dynamics assumed in the lattice model is approached when the local saturation time is much smaller than the typical time between long-range jumps; i.e. *τ*_s_ ≪ *τ*_j_. Using the above estimates for the characteristic times, we find the criterion

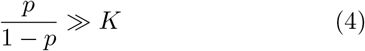

for fast local dynamics. This criterion is always satisfied as *p* → 1. When *K* is large, *p* must be at least 1 − 1*/K* for Eq. (4) to be satisfied: for appreciable local carrying capacities, the fraction of local dispersal events must be very close to one for the criterion to hold. If an individual competes with a large number of other individuals in its neighborhood for resources, Eq. (4) is satisfied only if the vast majority of dispersal events are local and long-range jumps are exceedingly rare. Our estimate emphasizes the need for simulations with explicit local dynamics to investigate the broad range of parameters where the lattice model assumptions do not hold during jump-driven range expansions.

The criterion *τ*_s_ ≪ *τ*_j_ ensures that new migrants originate from fully saturated regions. To satisfy the second assumption of the lattice model—the local founder-takes-all effect—we additionally require that the characteristic time between the arrival of the first migrant and a potential second migrant by long-range dispersal, which we call *τ*_2_, is much larger than the local saturation time scale *τ*_s_. Unlike *τ*_s_ and *τ*_j_, however, we do not have direct control over *τ*_2_; the latter time scale will depend not only on the model parameters but also on the location of the region being colonized. For example, the expected time to second arrival will be different for a region near the core of a colony compared with a region far from the core that was recently seeded by long-range dispersal. Nevertheless, we expect that *τ*_2_ is closely related to the time scale *τ*_j_ associated with sequential long-range jumps out of any given region: if long-range dispersal from all regions is exceedingly rare (*τ*_j_ is large), it will take a very long time for a second migrant to arrive into a newly colonized region (*τ*_2_ is large as well). Therefore, we use the same criterion, Eq. (4), to gauge whether both assumptions underlying the lattice model are satisfied in our continuum model. In the next section, we directly verify that our simulation results include regimes which violate the assumptions of instantaneous local saturation (Section IV A) and local founder-takes-all (Section IV B), thereby departing strongly from the prior lattice models. In summary, to violate the lattice model assumptions we require local dispersal probability values comparable to or lower than 1 − 1*/K*. If we evenly sample values of *p* between zero and one, we find that the lattice model assumptions are violated at most parameter values. For instance, if we set the carrying capacity to *K* = 10, the criterion is violated for *p* values up to around 0.9; when *K* = 100, the criterion is satisfied only for *p* > 0.99. In our simulations, we choose values of carrying capacity *K* between 10 and 100, and local dispersal probabilities in the range 0 ≤ *p* ≤ 0.997. We expect the lattice model assumptions to be violated over most of these parameter values, except at the upper range of values of *p*.

## IV. RESULTS

### A. Local dynamics are consistent with logistic growth

We first analyze the effect of modifying the local dispersal probability *p* on the population dynamics within interaction regions. Consider the fate of the interaction region surrounding a pioneer that has landed in an empty part of the range. If all local dispersal events experienced by the pioneer and its offspring landed within the pioneer’s interaction region, we would expect exponential growth of the local population with rate *p* until the carrying capacity *K* is reached. In practice, the interaction regions of the offspring only partially overlap with that of the pioneer, so the population growth levels off smoothly upon approaching the maximum value. When saturation curves across many interaction regions are averaged for a given set of parameters, the average curve takes on the form of a logistic function as shown in Fig. 4**a–b**. Upon varying *p* and *μ* independently, we find that the saturation proceeds faster as *p* is increased whereas it is not strongly affected by the kernel exponent (Fig. 4**b**).

**FIG. 4.**
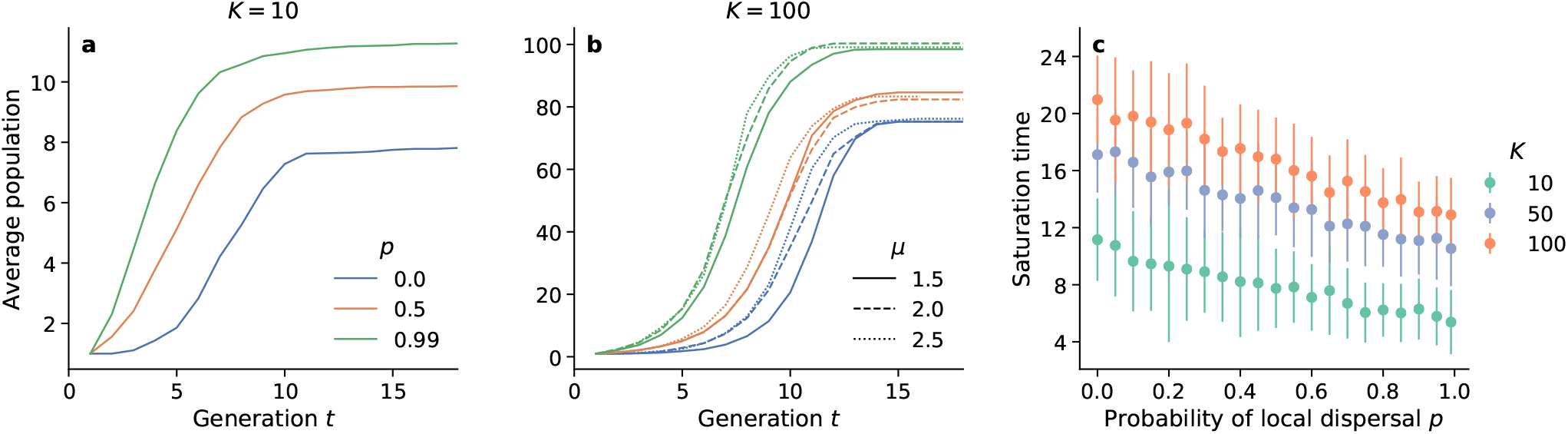
Saturation dynamics of interaction regions around pioneers. **a.)** and **b.)** show the population growth within the interaction region of pioneers (individuals which land in an empty region) as a function of time from establishment of the pioneer, averaged across many pioneers for different values of *p* (colors). **a**., *K* = 10 and *μ* = 1.5; **b**., *K* = 100 and three different kernel exponents (dashes). Each curve in panels (a.) and (b.) is the average of the local saturation around approximately 60 pioneers gathered across multiple simulations. **c.)** Saturation time of interaction regions, defined as the time taken for the fitted logistic growth function describing the population within an interaction region to reach one less than the saturating population (see SI Section VI C for details), for *μ* = 1.5. We fit the logistic growth function to the local saturation data of approximately 60 interaction regions around pioneers and then computed the saturation time for each region based on the fitted growth rate and carrying capacity. The points in the plot are averages and the error bars are the standard deviations of the computed saturation times of individual interaction regions at each set of parameters.

We use the logistic growth rate, extracted from a two parameter fit to the average growth curves (see SI Section VI C for details), to quantify the local saturation dynamics. As expected, we find that the growth rate is largely independent of carrying capacity and is determined by the local probability *p* (SI Fig. 12). The growth rate remains nonzero as *p* → 0, due to multi-step colonization: although no direct offspring of the pioneer can land in its own interaction region, the descendants of these offspring can land within the interaction region of the pioneer which eventually gets filled. Multi-step effects are also responsible for generating saturation curves whose final population values do not exactly equal the carrying capacity *K* (plateaus at large *t* in Fig. 4**a–b**), as outlined in SI Section VI C. The true saturation value of the population within an interaction region can be extracted from the logistic fit and is denoted as *K*′.

Although the logistic growth rate is set by the local dispersal probability and not the carrying capacity, the typical time taken to fill the interaction region of a pioneer depends on both quantities. Since the logistic growth function is continuous and strictly reaches *K*′ only as *t* → ∞, we define the time taken to reach a local population of *K* − 1 as the saturation time for an interaction region. We find that the saturation time falls with increasing local dispersal levels, and rises with increasing local carrying capacity, as shown in Fig. 4**c**. The interaction region around a pioneer that seeds a distant satellite takes longer to fill up at low local dispersal rates and/or at high carrying capacities. Notably, the saturation time falls linearly with *p*, but has a slow (roughly logarithmic) functional dependence on the carrying capacity.

### B. Slow local saturation invalidates founder-takes-all assumption within interaction regions

Slow saturation of the pioneer’s local region increases the chance that other individuals who are not descendants of the original pioneer will disperse into the region and establish themselves before the region is full. If an individual who arrives later has a different allele than the original pioneer, there will be multiple alleles within the region, which introduces genetic diversity within interaction regions in stark contrast to the homogeneous demes imposed by the lattice model. This creates a measurable signal that local saturation times are now comparable to or slower than the typical time gap between the arrivals of first and second migrants by long-range dispersal, *τ*_2_.

To quantify the deviation of local population structure from the local founder-takes-all assumption as the saturation time is increased, we introduced neutral genetic variation in the initial population. Every individual in the initial population was assigned a unique allele, which did not affect the dispersal or reproduction dynamics but was passed on to offspring. The establishment of multiple alleles in the same interaction region was detected by computing the local heterozygosities in the interaction region of isolated pioneers. The heterozygosity, *H*, is the probability that any two randomly selected individuals will have different alleles. Upon counting the fraction *f*_*i*_ of individuals with each neutral allele *i* in an interaction region, the heterozygosity of that region is computed as

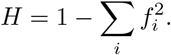

A nonzero heterozygosity indicates that more than one allele is present in the region; the larger the heterozygosity, the more evenly distributed the different alleles are in frequency, corresponding to a region in which no single allele dominates.

We averaged the local heterozygosity within the interaction regions of many independent pioneers to obtain a characteristic measurement of the local diversity for each parameter value. The averaged heterozygosities are normalized against the value at which one expects a fully occupied interaction region to have exactly one individual with a different allele than the pioneer: *H*_N_ ≡ 2(1*/K*)(1 − 1*/K*). With this definition, the normalized average heterozygosity ⟨*H*⟩ */H*_N_ has the following interpretation: normalized average heterozygosities less than one indicate that interaction regions typically have a single allele, whereas values greater than one indicate the expected presence of more than one allele signaling a deviation from the founder “taking all” at the level of the interaction region.

We find that the local heterozygosity is high at low local dispersal rates and high carrying capacities (Fig. 5), consistent with our expectations from the slow saturation dynamics in this part of parameter space. At the smallest carrying capacity (*K* = 10), heterozygosity levels are low across nearly all jump kernels: local saturation occurs fast enough that interaction regions are filled by descendants of the pioneer individual that first arrived in the vicinity. This situation most closely parallels the lattice models. As the carrying capacity is increased, however, we observe appreciable levels of heterozygosity at low levels of local dispersal where the saturation dynamics of regulation regions is slowest (Fig. 4c). As the local dispersal rate increases, a smooth crossover occurs from high to low heterozygosity. The value of *p* at which this crossover occurs is larger for broader jump kernels (lower *μ*): longer dispersal events favor mixing of alleles. We expect these trends to continue for carrying capacities on either side of the range we show here. For lower carrying capacities, local diversity would become lower everywhere. For higher carrying capacities, the boundary between pioneer-dominated and not pioneer-dominated (blue points and red points, respectively) would continue to move to the right. The region of parameter space where founders typically “take all” will continue to shrink as carrying capacity increases.

**FIG. 5.**
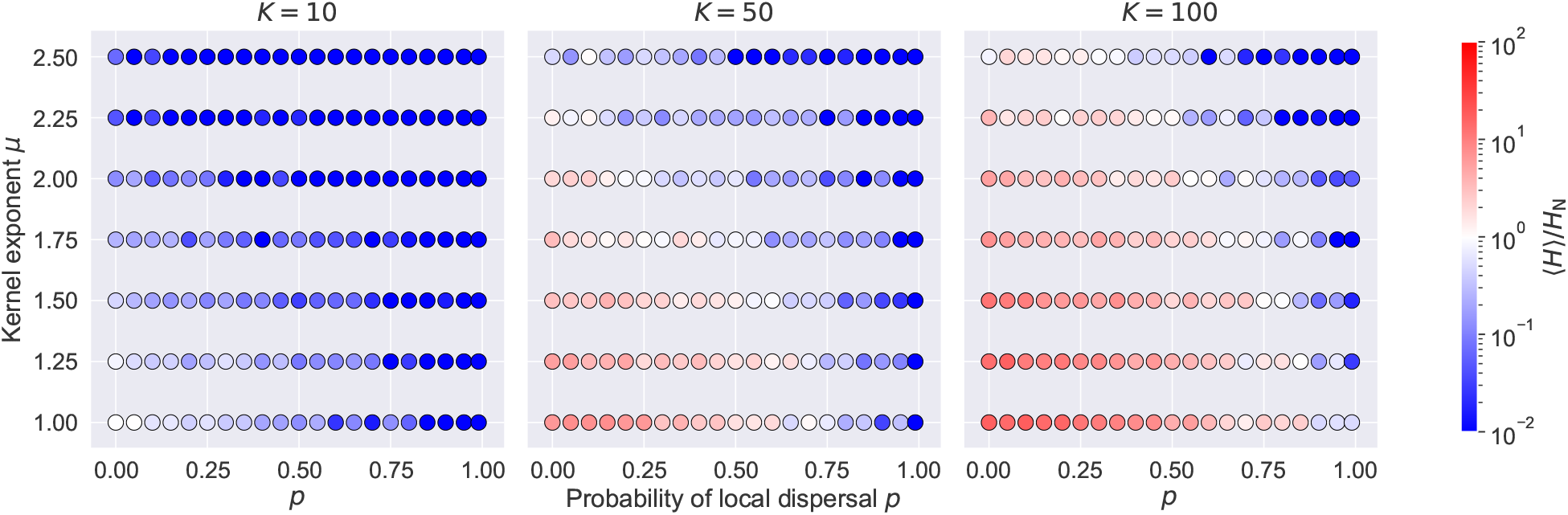
Influence of slow local dynamics on local diversity. Each symbol represents a triplet of parameter values (*K, p, μ*) and its color shows the average heterozygosity within the interaction regions around several pioneers who seeded distant satellites, normalized against the heterozygosity at which a fully occupied interaction region is expected to have one individual with an allele different from everyone else in the region. Interaction regions are expected to be homogeneous at parameter combinations where the average normalized heterozygosity is less than one (blue points). They are expected to have more than one allele where the average normalized heterozygosity is greater than one (red points), indicating that other individuals typically disperse into and establish themselves within a pioneer’s interaction region before it fills up with descendants of the pioneer. The values reported come from the averages across about 50 interaction regions gathered from multiple simulations at each set of parameters.

In summary, measurements of local heterozygosity (Fig. 5) indicate a breakdown of founder takes all over wide swaths of parameter space, especially for high carrying capacities and broad jump kernels. While local interaction regions remain largely monoallelic when long-range dispersal is very rare (*p* ≳ 0.9), we find evidence that multiple incursions into the same region leave a persistent contribution to the local genetic makeup within interaction regions when long-range and local dispersal rates are of similar order. We next investigate the extent to which these *local* deviations from founder takes all impact *global* features of the population expansion, and in particular whether they lead to departures from the population-level behavior of jump-driven range expansions predicted using lattice-based models in Refs. [16, 27].

### C. Increased long-range dispersal favors faster population growth

The salient feature of the global population growth under jump-driven expansions is their dramatic speedup compared to expansions that only involve short-range jumps: the typical radial extent of the core region ℓ(*t*) grows faster-than-linearly with time when *μ* < *d* + 1. This boost occurs because offspring attempting shortrange jumps will land close to their parents and siblings, and are more likely to be unsuccessful due to a lack of local carrying capacity. By contrast, long-range jumps tend to transport offspring to empty areas where they establish and proliferate successfully. Therefore, lower values of the local dispersal probability *p* are expected to favor faster population growth overall, even though the local saturation is slower.

We measured the population growth with time, *M* (*t*), for many independent range expansions at each parameter value. To connect with the results from lattice-based models described in Section II A, we need an estimate of the core region within which the population has reached saturation. When growth is driven by long-range jumps, there is no sharp boundary between occupied and empty regions even in the lattice model. Rather, the local density is close to *ρ* out to some distance from the origin, beyond which it crosses over to a power-law decline in density determined by the value of *μ* [16, 39]. This smoothly varying occupancy profile leaves some ambiguity in precisely defining the core region. We follow Ref. 39 in using the mass-equivalent radius of the population as our best estimate of the core radius from our simulation data:

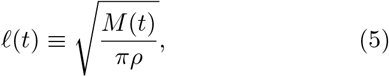

which provides the required scaling *M* (*t*) ∼ [ℓ(*t*)]^*d*^ [16]. This definition assumes that the bulk of the population is present in regions where the population has reached its maximum density locally. We averaged ℓ(*t*) trajectories across different instances at each set of parameters to get a *growth curve* characterizing the average growth in extent of the population.

We found that the acceleration of range expansion due to long-range dispersal is preserved in the continuum model, as shown by the growth curves in Fig. 6. We focus on the behavior at long times beyond the saturation time scale of a single interaction region (which is of order 10 for *K* = 10, see Fig. 4**c**). When all dispersal is short-range (*p* = 1), the average colony size approaches a linear relationship at long times (dashed curves; linear fit shown with solid curves at upper right), signifying the expected constant-speed outward advance of the population front [30]. Small levels of long-range dispersal (solid curves) are sufficient for the size to grow faster than linearly with time, as evidenced by a steeper slope on log-log axes compared to the dashed curves. The growth at long times appears to be faster than any power law (i.e. faster than linear on log-log axes) for all values of *p* at *μ* = 1.5 and *μ* = 2.0 (Fig. 6**a–b**), in line with expectations from the lattice model. Growth approaches a power law in time with exponent greater than one at the the two largest *p* values for *μ* = 2.5, but is faster than power-law for the smaller values of *p* over the population sizes simulated. In all cases, decreasing the probability of short-range dispersal speeds up the colony expansion, as expected: long-range jumps are far more likely to land in empty regions and succeed, compared to local jumps.

**FIG. 6.**
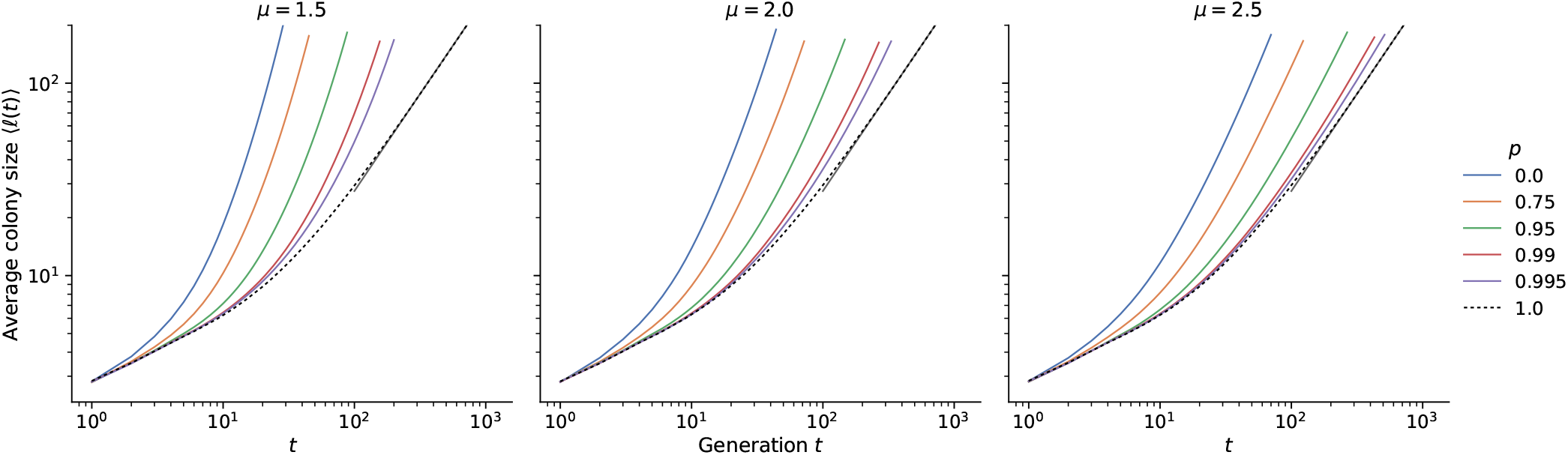
Average growth curves for different *μ* and *p* at *K* = 10. These are averages of growth curves from 241 individual simulations at each set of parameters. Average curves at *K* = 100 are shown in Fig. 11. The dashed line corresponds to simulations with only local dispersal; the resulting growth at long times is consistent with a linear relationship (gray line in the upper right of each panel).

Many consequential features of the expansion, however, are determined not by the absolute growth of the population size with time but by the functional form of the growth. For instance, the qualitative differences in global diversity among different kernel ranges (Section II B) are owed to the different functional forms of ℓ(*t*) observed in the lattice model (see Section II for a summary). It would be useful to quantify whether and how the local dispersal rate influences the functional form of the population growth curves. A direct comparison of the growth curves to the asymptotic forms derived using the lattice model is not expected to succeed, because the growth curves can take a long time to reach their asymptotic forms, especially for values of *μ* near the space dimension *d* = 2 [16]. This feature of jump-driven growth is apparent in Fig. 6**c**, in which the measured growth curves for *μ* = 2.5 are nonlinear on logarithmic axes and deviate from the asymptotic power-law form even at long times. Instead, we use the self-consistency condition from Ref. 16, Eq. (1), which is expected to hold for times beyond the local saturation time scale but well before the time at which the asymptotic regime is reached in the lattice model. If the entire population in our continuum model were contained in regions that have reached local saturation at all times, then the hierarchy depicted in Fig. 1 would translate to the continuum model as well, and we would expect Eq. (1) to be satisfied exactly. This would enable us to predict future population growth given only the current population size and the exponent that characterizes the jump kernel. The size of deviations from the exact relation could be used to quantify differences in satellite structure between the continuum model and the lattice model.

To test the validity of the consistency condition and its ability to predict population growth, we measured the relationship between the colony size ℓ(*t*) at time *t* and the quantity *t*ℓ(*t/*2)^2*d*^ in our simulations. For *t* values larger than the local saturation time (order 20 or less for all parameters, Fig. 4**c**), we found that the simulated growth curves are consistent with a power-law relationship between the two quantities across the entire range of local dispersal probability values tested. Data for two representative values of *p* and two local carrying capacities are shown in Fig. 7; additional curves are shown in SI Fig. 14. For parameter values which best approximate the assumption of instantaneous filling of density regulation regions (local dispersal probability close to one and low carrying capacity), the power-law exponent quantifying the relationship between ℓ(*t*) and *t*ℓ(*t/*2)^2*d*^ also matches the expected exponent of *d* + *μ* (compare green discs to dashed line in Fig. 7). By contrast, the relationship no longer quantitatively matches the consistency condition when local saturation is slowed down by low values of *p* or high values of *K*. Instead, the population size at time *t* is larger than that predicted by the population at time *t/*2 according to Eq. (1) (square symbols and orange discs in Fig. 7). The functional form of the growth curve appears to be *faster* than would be expected from the time doubling hierarchy, so using Eq. (1) leads to an underestimate of the colony size at time *t* given its size at time *t/*2. Note that Fig. 7 is plotted on logarithmic axes, so the visibly slight difference between the slopes of sets of symbols and dashed lines corresponds to different power law relationships between ℓ(*t*) and *t*ℓ(*t/*2)^2*d*^ in our continuum expansions than what is predicted by the time doubling hierarchy derived using lattice models that assume instant saturation of local regions.

**FIG. 7.**
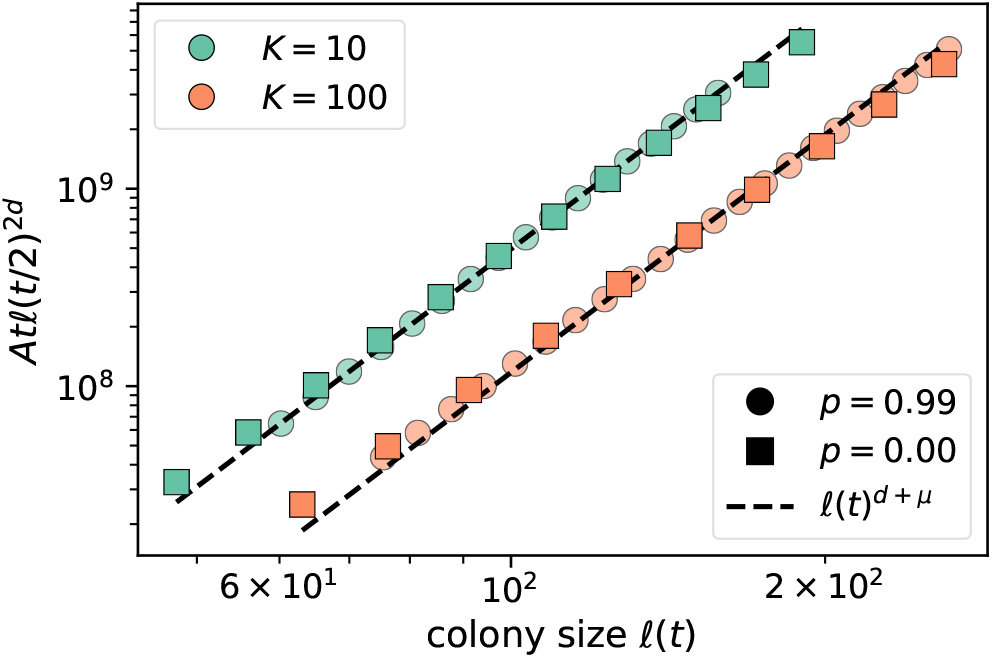
Quantitative test of the hierarchical time-doubling structure. Plots show the RHS of the consistency condition (Eq. (1)) versus the colony size ℓ(*t*) for *μ* = 2. Data are from the average of about 200 growth curves at each set of parameters, and only include the second half of the simulation to exclude expected deviations at short times (see SI Section VI D for details). The scaling factor *A* was adjusted manually to overlay data from different *p* values for ease of comparison of the apparent power-law exponent (slope of curves on log-log scale). Analogous plots at *μ* = 1.5 and *μ* = 2.5 are shown in SI Fig. 14.

To quantify the extent of the deviation from the lattice-model behavior, we fit measurements of the quantity *t*ℓ(*t/*2)^2*d*^ to the form *B*ℓ(*t*)^*ν*^ to extract the power-law exponent *ν* (see SI Section VI D for details). This exponent was used to infer a kernel exponent *μ*_i_ ≡ *ν* − *d* from data such as those shown in Fig. 7, which can be compared to the true kernel exponent *μ*. To cover the two distinct jump-driven growth regimes and the marginal value *μ* = *d* separating them (as referenced in Section II), we estimate the kernel exponent from growth curves of populations whose jump kernels decay with *μ* equal to 1.5, 2.0, and 2.5. We find that the inferred kernel exponent is close to the true exponent when the local dispersal probability approaches one across all jump kernels and carrying capacities tested (Fig. 8). This observation is consistent with our expectation that the limit *p* → 1 best approximates the lattice model assumptions. However, the inferred kernel exponent is systematically lower than the true value for much of the range 0 < *p* < 1, reflecting the shallower-than-expected slopes at low local dispersal in Fig. 7. The inferred exponent grows slowly with the local dispersal probability up to *p* ≈ 0.9, and then rises sharply toward the true value as *p* → 1. This suggests that there could be some functional change to the structure of colony expansion as the parameter changes to nearly all short-range dispersal, while anything less than nearly all short-range dispersal seems to behave similarly regardless of *p*. The deviations also systematically differ depending on the local carrying capacity, with inferred exponents at *K* = 100 consistently lower than than those at *K* = 10.

**FIG. 8.**
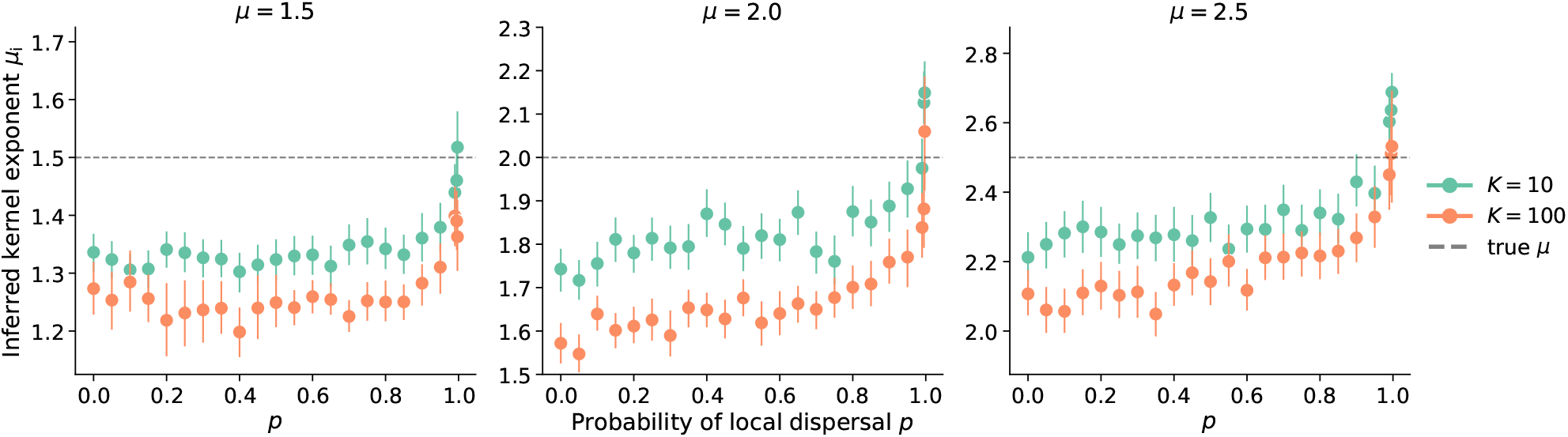
Inferring kernel exponents using the consistency condition. Symbols show the kernel exponent *μ*_i_ inferred for the power law relationship between measured ℓ(*t*) and *t*ℓ(*t/*2)^2*d*^ (i.e. the slope in Fig. 7) from many individual simulations. Panels are labeled by the true kernel exponent *μ* used in simulations. The dashed line indicates *μ*_i_ = *μ*. Each point represents the mean of the individual inferences from roughly 200 independent simulations and the error bars are the 95% confidence interval of the distribution of bootstrapped mean inferred kernel exponents.

We have not isolated the mechanism leading to an inferred kernel exponent *μ*_i_ that deviates from the true kernel exponent *μ*. The fact that *μ*_i_ < *μ* implies that the time-doubling hierarchy from the lattice model, quantified in Eq. (1), does not hold exactly over much of the range of *p* values. Furthermore, it shows again that the functional form of the population growth with time is faster in the continuum model than the lattice model. However, this observation by itself does not provide information about how the hierarchy breaks down in the continuum model, or whether a modified version of Eq. (1) might be found for continuum space models.

We can nevertheless identify the likely sources of the discrepancy between the continuum and lattice models based on our knowledge of the local and global dynamics. The hierarchy in the lattice model was derived under the assumption that satellites which drive the expansion originate in a core region that has reached its saturation density nearly everywhere, and whose size scales as [*M* (*t*)]^1*/d*^. In our simulations, local regions take some finite amount of time to fill up, but they can begin sending out long range migrants as soon as they are seeded. An appreciable fraction of satellites may be seeded by individuals dispersing from regions with local densities between zero and *ρ*; furthermore, the local density could itself vary significantly through the population. These deviations become more prevalent for larger carrying capacities (Fig. 4**c**), which would suggest larger deviations at higher values of *K* consistent with the behavior of the inferred kernel exponents in Fig. 8.

Altogether, measurements in the continuous-space model reveal small but consistent deviations in the population growth curves from the time-doubling hierarchy predicted in the lattice model. Our simulations indicate that slow local dynamics introduce corrections to the time-doubling hierarchy over a large range of values of the local dispersal probability, consistent with our estimates of the parameter regimes for which the lattice model assumptions break down (Section III). Next, we numerically investigate the impact of these corrections on the dynamics of global diversity, for which the hierarchy of satellite sizes determined the long-time behavior in the lattice model as summarized in Section II B.

### D. Increased local diversity boosts global heterozygosity but does not overcome long-term trends

Finally, we investigated the consequences of the enhanced local diversity generated by slow local dynamics (Section IV B and Fig. 5) on the fate of the initial neutral diversity. Recall the predicted evolution of heterozygosity in prior models assuming fast local saturation and local founder-takes-all effects (summarized in Section II B): initial variation decayed steadily towards zero for jump kernels with 2 < *μ* < 3 in two dimensions, but some proportion of the initial diversity was preserved for broader (*μ* < 2) or narrower (*μ* > 3) kernels. We simulated range expansions where the initial population had equal proportions of two fitness-equivalent alleles (initial global heterozygosity *H*_G_ = 0.5) and measured the evolution of global heterozygosity. While the outcome of a single simulation is stochastic, we estimated the expected value of the hetorozygosity as a function of population size by averaging the outcomes of many independent runs at each set of parameters. Recall that no new mutations appear during the expansions; here we study the long term fate of any pre-existing diversity present in the initial population rather than the emergence of some balance between the loss of diversity (e.g. due to drift) and the promotion of diversity due to new mutations. Although we used a specific initial heterozygosity in our simulations, we expect the observed trends in the *proportion* of initial diversity over time to hold for other values of initial global heterozygosity as well. This proportion is obtained from our simulation data (Fig. 9) by dividing the reported heterozygosity values by 0.5.

**FIG. 9.**
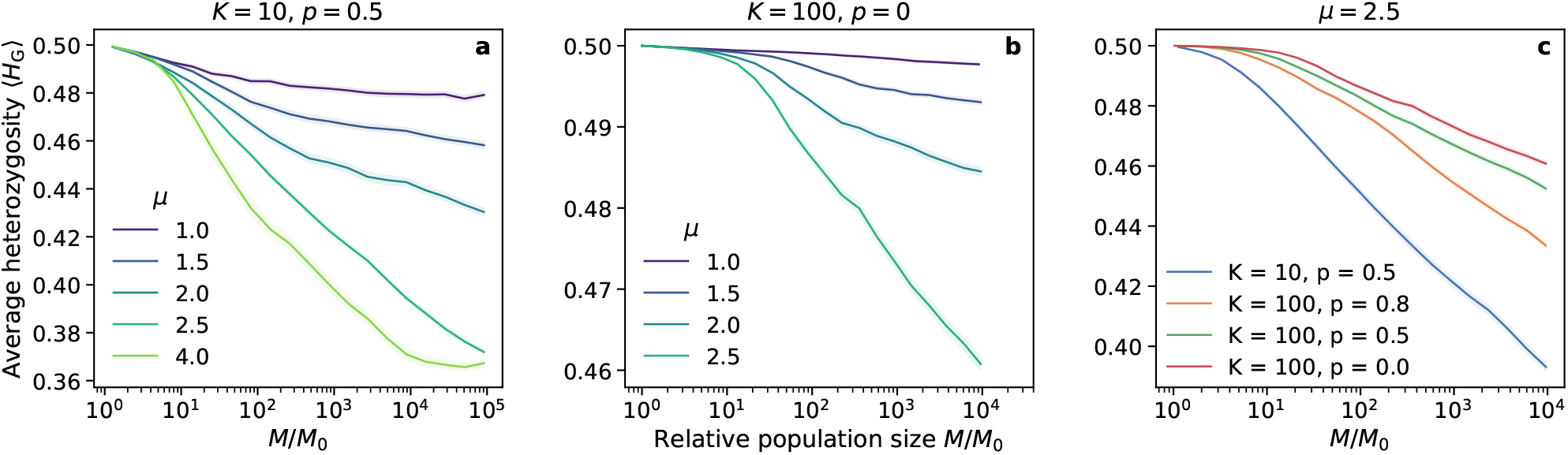
Evolution of global heterozygosity for different kernels and local dispersal rates. **a.)** Average global heterozygosity as a function of the growing population size for different jump kernels, with *K* = 10 and *p* = 0.5. At these parameters, each interaction region is dominated by a single allele (Fig. 5). **b.)** Same as **a** for *K* = 100 and *p* = 0; at these parameters interaction regions tend to harbor multiple alleles. **c.)** Global heterozygosity curves for *μ* = 2.5 and different local dynamics traversing the spectrum from monoallelic to multiallelic local interaction regions (blue points to red points in Fig. 5). The *K* = 10 data is the same as in panel (a.) but has been truncated for this plot. Data as a function of relative population size were generated by by binning the population sizes from all available simulations and then computing the within-bin ⟨*H*_G_⟩ ; see SI Section VI E for details. Shading reports the standard error of the mean within each bin in all panels, as an estimate of the uncertainty in our estimate of the ensemble average. Data come from about 200 independent simulations for *μ* ≤ 2 in panel (a.) and about 400 simulations for *μ* = 2.5 in panel (a.) and all of panels (b.) and (c.). The curve for *μ* = 4 in panel (a.) comes from just 24 runs since simulations with *μ* > *d* + 1 take much longer and the sectoring mechanism for preserving diversity is well understood (see Fig. 2**a–c**).

We first considered a set of parameters (*K* = 10, *p* = 0.5) for which each interaction region is dominated by the offspring of the seed individual (Fig. 9**a**). This situation approximates the local founder-takes-all mechanism of the lattice models, but does not replicate it exactly as multiple incursions into interaction regions are not strictly excluded. Despite the deviations, we find that the evolution of global diversity in the continuum simulations is consistent with expectations from the lattice model when different kernel exponents are compared. (See SI Section VI F for a quantitative comparison.) Average population heterozygosity has settled to a constant proportion of its initial value for *μ* = 1, and appears to be approaching a constant value as well for *μ* = 1.5. The slow decay of heterozygosity for *μ* = 2 is expected; the population may have to grow by several more orders of magnitude before converging to a constant heterozygosity [27]. At *μ* = 2.5, the heterozygosity decays steadily with no sign of convergence to a finite value, as predicted for lattice models in the range *d* < *μ* < *d* + 1. At *μ* = 4, a constant heterozygosity is attained at large population sizes due to the formation of persistent sectors with distinct allelic identities (Fig. 2**a–c**). In each of the growth regimes separated by the critical kernel exponent values of *d* and *d*+1 (2 and 3 respectively in our two-dimensional expansions), the behavior of the global heterozygosity follows the qualitative patterns derived in the lattice model. In spite of the small quantitative differences in the hierarchical structure of satellites merging with the core (Fig. 8), the overall differences in structure which determine the balance between diversification and coarsening in jump-driven expansions are maintained deep within the different growth regimes.

Next, we considered parameters *K* = 100, *p* = 0 for which the local founder-takes-all assumption is violated across all kernels tested according to local heterozygosity measurements. We found that the increased local diversity at these values (as indicated by colors in Fig. 5**c**) contributes to higher global heterozygosities compared to the fast saturation region, as seen in Fig. 9**b** when compared to Fig. 9**a** and SI Fig. 16. For instance, at *μ* = 2.5 the heterozygosity has decayed by around 8% of its initial value when *M/M*_0_ = 10^4^ in Fig. 9**b**, in contrast to a reduction by over 20% in Fig. 9**a**. The same trend is observed at all kernel exponents: The mix of allelic identities within each interaction region under slower local dynamics provides a reservoir of genetic diversity that allows populations to retain much more diversity than possible under the monoallelic regions imposed by fast local saturation. Nevertheless, the qualitative trends in diversity as the kernel exponent is varied continue to track the expectation for lattice models. In particular, the global heterozygosity steadily decays towards zero for *μ* = 2.5, albeit at a slower rate compared to the *K* = 10 simulations.

A steady decay in heterozygosity is also observed for other values of the local dispersal probability for the same kernel exponent *μ* = 2.5, see Fig. 9**c**. Slowing down local dynamics by increasing *K* and reducing *p* raises the value of heterozygosity at each population size, but does not prevent the steady decay as a function of *M/M*_0_. These results show that at long times, the diversity-reducing effect of bottlenecks outweighs the local mixing due to slow saturation dynamics for *μ* = 2.5. We expect that continual heterozygosity loss will be experienced for other kernels in the range *d* < *μ* < *d* + 1 as well, although the rate of decay will be very slow for kernels close to the critical value of *μ* = *d*, and for kernels with slower local dynamics (i.e. large carrying capacity and low local dispersal probability). In this regard, the high local heterozygosities observed for kernels with *μ* > 2 and low *p* values in Fig. 5 are transients which we expect to decay to lower values if the expansions are allowed to run longer.

## V. DISCUSSION

Range expansions in populations experiencing long-range dispersal can be dominated by the pioneers who travel long distances and seed satellite colonies. Lattice models that assume that these pioneers quickly saturate the carrying capacity within their local interaction region have provided many insights into the dynamics and population structure of such range expansions [16, 27]. However, real populations operate in continuous space and with local population dynamics which play out concurrently with the global dynamics driven by long jumps. In particular, the limits on the rates of long-range dispersal for lattice models to be accurate become increasingly strict as the local carrying capacity increases (Eq. (4)). We have introduced a continuous-space simulation of range expansions which departs from the gridlike spatial structure and instantaneous local dynamics implied in lattice models, enabling us to quantitatively investigate population growth and neutral diversity in parameter regimes where the lattice models are not expected to be valid.

We found that introducing explicit local dynamics is associated with slow local saturation at low local dispersal rates and especially at high local carrying capacities (Fig. 4). By contrast, the global population growth occurs faster when local dispersal rates are low, because of the increase in long-range jumps that seed satellite populations in unoccupied regions (Fig. 6). The functional forms of the population growth curves show similarities with those from lattice-based models (Fig. 7), but with small yet quantifiable differences (Fig. 8). We suspect that these differences arise due to a violation of a central assumption of the lattice model: that satellites are seeded by long-range migrants who disperse from *fully occupied* source regions. In our continuum model, satellites can begin sending out long-range migrants as soon as they are seeded, which can occur several generations before they saturate at high carrying capacities and low local dispersal rates. In future work, we aim to incorporate this feature into the model of hierarchical population growth sketched in Fig. 1, which would improve the accuracy of theoretical predictions for jump-driven range expansions in situations where local interaction regions are not immediately saturated upon the arrival of a new migrant.

We investigated the effects of departing from instantaneous local saturation on both local and global measurements of neutral diversity. Interrogating the populations within individual interaction regions originally seeded by a long-range dispersal event reveals that multiple lineages, rather than just descendants of the pioneer, become likely as local saturation becomes slower (Fig. 5): our continuum model violates the assumption of a strictly enforced local founder-takes-all effect. Having multiple lineages within interaction regions provides a reservoir of genetic diversity that also enables greater global heterozygosity outside the regime where local founder-takes-all applies: generically, expansions with slower local dynamics exhibit higher global diversity at every stage in the expansion (Fig. 9). Nevertheless, the enhancement in local diversity is not sufficient to overcome long-time trends in global diversity, which continue to be determined by the kernel exponent as was shown in the lattice model [27]. In particular, when *μ* < 2 the global heterozygosity settles to a stable value after an initial period of decay, whereas for 2 < *μ* < 3 the heterozygosity decays steadily as the range expansion progresses albeit at a slow rate. The decay is a consequence of the repeated coarsening of diversity due to bottlenecks as pioneers expand into their newly occupied surroundings (Fig. 2**d–f**). Our results show that this coarsening is slowed down by the increased local diversity when the local founder-takes-all assumption is violated, but it is not completely mitigated and the qualitative long-term trends in global diversity are similar to those predicted using the lattice model. This qualitative agreement with lattice-based predictions is a non-trivial result in light of recent research [35] showing that models based on a discretization of space can leave surprising artifacts in measures of population genetic variation.

Our method of discovering local diversity outside the local founder-takes-all regime was unable to detect if descendants of an individual other than the pioneer were within a local region if they happened to have the same allele as the pioneer by chance. Such information would be useful to investigate genealogical structure beyond the fate of the initial neutral diversity in the population, for example to determine if the pioneer is the most recent common ancestor of everyone else in the interaction region or to study the accumulation of additional neutral mutations during the expansion. A tool like tree sequences [41] could readily be incorporated into our computational model to study such questions, which are a promising target of future work. Understanding the competing effects of local and long-range dynamics on genealogies in our forward-in-time simulations could also aid the construction of backward-in-time models that incorporate long-range dispersal [42, 43].

Another promising future direction would be to incorporate ongoing local competition among all individuals in the population. In this work, we assumed that established individuals never move or die, modeling populations such as trees which release large numbers of seeds annually and where young saplings stand little chance of outcompeting mature trees around them. However, there are many species of perennial plants, for example, where younger individuals can successfully compete against older individuals in their surroundings. Incorporating population renewal and density-dependent competition in simulations could provide new insights into how these species evolve during range expansions. We suspect that such competition should accelerate the decay of diversity relative to our results for 2 < *μ* < 3 (Fig. 9). Local competition can completely remove alleles from the population, whereas in our model the “losing” allele is surrounded but not lost, and retains a nonzero probability of dispersing an offspring to a faraway vacant habitat.

This work provides a better understanding of the range of validity and the limitations of models of long-range dispersal which rely on instantaneous saturation of local interaction regions and divide continuous space into a lattice. We have confirmed that the conclusions of the lattice model are upheld in populations where pioneers who disperse long distances quickly saturate their immediate surroundings with their descendants; namely in populations with low local carrying capacities and high local dispersal probabilities. Even when the local founder-takes-all condition is violated, we have shown that qualitative trends in population growth and in the evolution of neutral diversity mirror those in the lattice model, albeit with measurable quantitative differences. Heuristics such as the time-doubling hierarchy of Ref. 16 (Fig. 1) and the effective population of satellites identified in Ref. 27 remain useful to understand the qualitative behavior of expansions under long-range dispersal in non-lattice models. Researchers could employ hybrid discrete/continuous research strategies: identify regimes of interest using the heuristics of the lattice model, and then test and refine these predictions in more realistic continuum simulations.

Our results are relevant to understanding and modeling the dynamics of range expansions in true biological populations, including invasive species, populations fleeing climate catastrophes, and spreading viruses. We now have a better understanding of when the first individual to arrive in a region of space effectively determines the genetic outcome of all others who will later inhabit the same immediate area. Experimenters could estimate the size of the interaction region, the local carrying capacity, and the local dispersal probability in populations of organisms in the lab or in nature. Estimates of those quantities could allow researchers to predict whether or not the founders will “take all” when the population expands its range outwards into new territory, leading to insights about how the population will evolve.

## DATA AVAILABILITY

Simulation code and the code and data necessary to generate figures are available in the following GitHub repository: https://github.com/paulose-group/explicitlocal-dynamics

## ACKNOWLEDGMENTS

We thank Peter Ralph for helpful comments on a draft of this paper. This work benefited from access to the University of Oregon high performance computing cluster, Talapas.

## VI. SUPPLEMENTARY INFORMATION

### A. Simulation details

Simulations begin with 10*K* individuals who are given random positions near the origin. Their *x* and *y* positions are random draws from a normal distribution with a standard deviation of 2*r*_b_. Typically about 80% of those individuals survive the density regulation in the first time step. The spatial landscape is large, so the periodic boundary conditions have no effect.

Offspring are produced by cloning without the possibility of mutations, so offspring have the same allele as their parent. Dispersal distances are drawn using inverse transform sampling. Recall, the jump kernel is

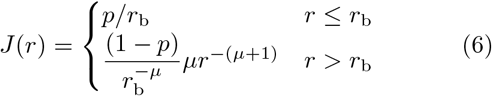

where *r*_b_ is the boundary between local and long-range and *p* is the probability of dispersing within the local region.

We begin the sampling procedure by drawing a random number *X* from the uniform distribution between 0 and 1. That number *X* is taken to be the the probability of drawing a dispersal distance less than or equal to some distance *x* (i.e. the integral of the jump kernel from 0 to *x*). Solving for *x* gives us our dispersal distance.

If *X* ≤ *p*, the offspring disperses locally, so we only need to consider the first term of the jump kernel.

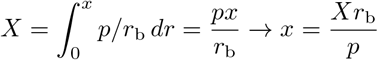

If *X* > *p*, the offspring disperses a long distance, so we have

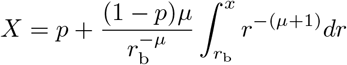

which leads to

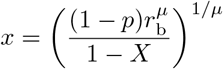

The dispersal direction is chosen at random from the uniform distribution between 0 and 2*π*.

All individuals in the population get a chance to produce offspring each time step. Offspring generation is the first thing that happens each time step; the number of offspring for each individual is a random draw from the Poisson distribution with mean 1. Then all newborns *simultaneously* count how many other individuals are within their density regulation regions. This means newborns will count other newborns if they happen to land near each other by chance. It also means that space often doesn’t quite fill up to the population density 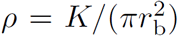 (as defined in section III). SI Fig. 10 shows how regions of space may appear habitable, and indeed would be if one single offspring were generated and counted its neighbors at a time, but do not saturate since everyone in the population typically produces an offspring every generation and all newborns count their neighbors simultaneously. All individuals produce one offspring per generation on average, so a region saturated to *K* individuals is expected to have roughly *K* newborn individuals attempting to establish within that same region every generation. Our density regulation mechanism mimics the biological scenario where none of those newborn individuals are able to get enough resources to survive since there are so many competing for what little is left, a situation termed “scramble competition” in ecology [44]. The typical population that is actually attained in a local density region, which we term *K*′, is estimated using a fit to a logistic growth curve (see SI Section VI C below), and deviates by at most 20% from *K* (SI Fig. 12). Alternative choices for the density regulation step, such as randomly choosing a subset of newborns to survive so that local density regions can saturate up to the target value *K* (the “contest competition” scenario), could also be implemented, but at the cost of additional computational resources which would affect the maximum population sizes and growth times that could be simulated.

**FIG. 10.**
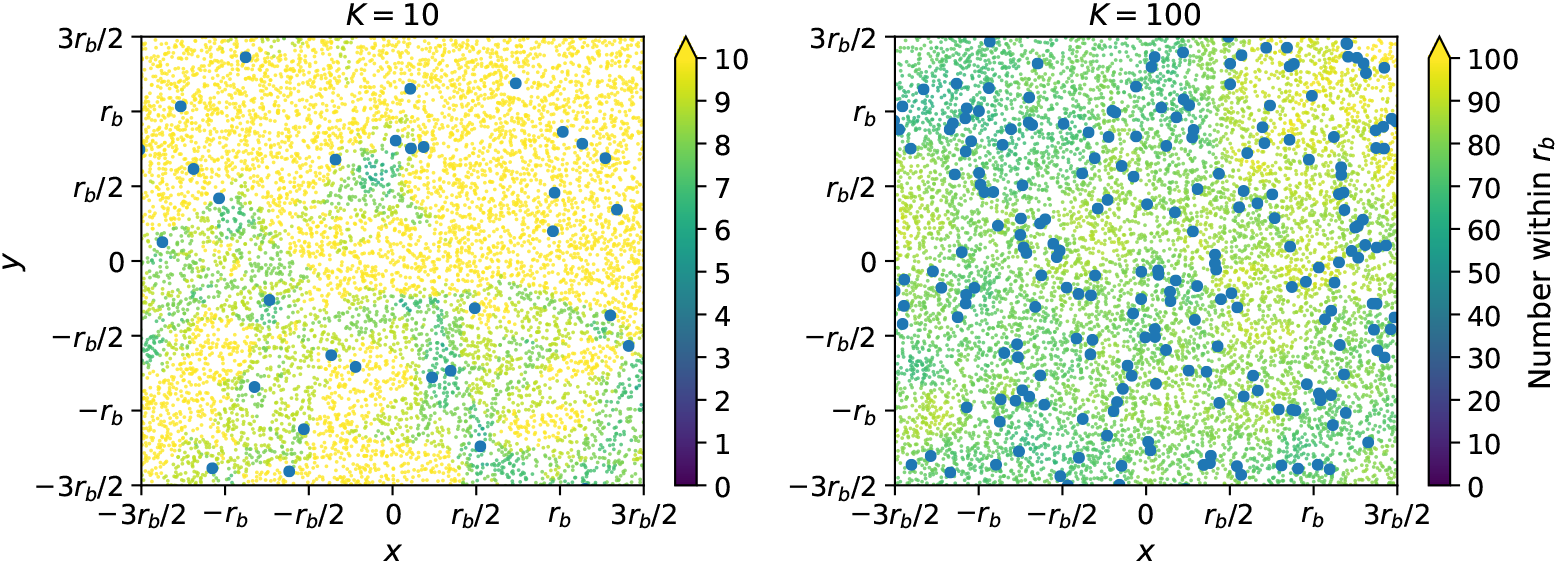
Space doesn’t quite fill up to the local maximum population density *ρ*. The two panels are snapshots from simulations that grew to roughly fifty thousand individuals. The blue points are individuals, and the color of the small yellow and green dots represents how many individuals are within a distance of *r*_b_ of that point. Yellow points represent saturated regions; an individual born there would count at least *K* within its density regulation region. Non-yellow points look hospitable, and one individual born there would count less than *K* within its density regulation region, but individuals can’t fill those spaces since typically everyone produces an offspring every generation and the newborns “destructively interfere”.

We typically let the populations grow by about four orders of magnitude, so simulations were ended once the populations exceeded 10^6^ or 10^7^ individuals for *K* equal to 10 or 100, respectively. This allowed core radii to grow by about two orders of magnitude, as shown in the average growth curves at *K* = 10 in Fig. 6 and at *K* = 100 in SI Fig. 11. The solid line indicating the linear relationship between ⟨ℓ(*t*) ⟩ and *t* at *p* = 1 in Fig. 6 was generated by fitting a line to the average growth curve from generations 100 to 1000 using NumPy’s polyfit() function.

**FIG. 11.**
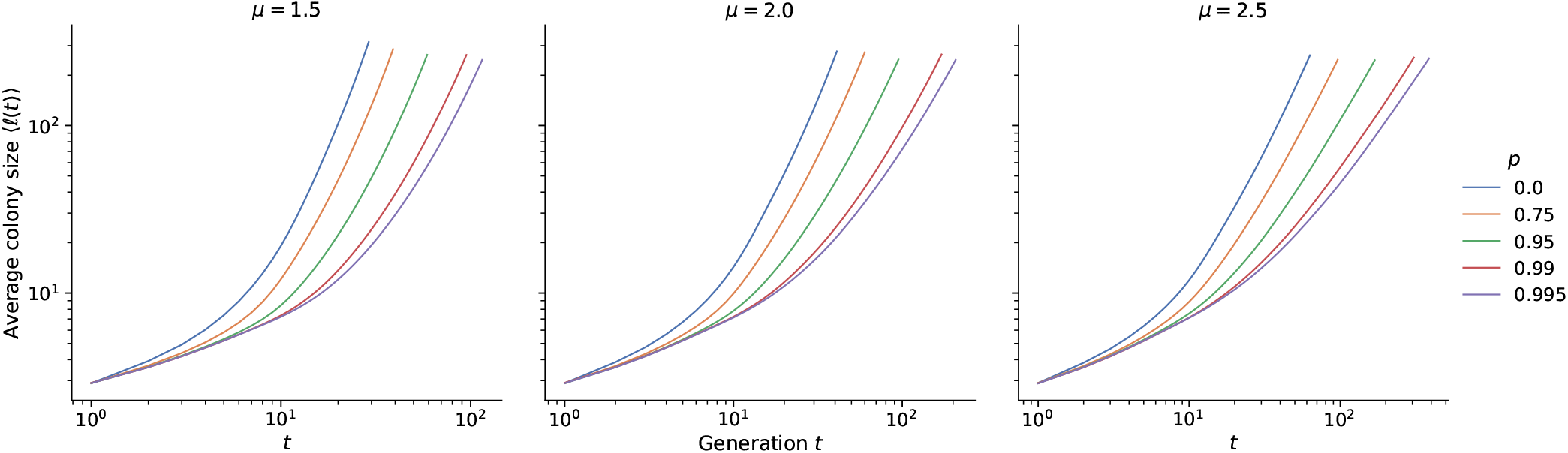
Average growth curves for different *μ* and *p* at *K* = 100. Average curves for *K* = 10 are shown in Fig. 6. These are averages of about 140 growth curves at each set of parameters.

### B. Time scales

The assumption of instant local saturation in the lattice models relied on a separation of time scales between local and global dynamics: it is valid provided the time scale for saturation of local regions *τ*_s_ = 1*/α* is small compared to the typical time between long-range dispersal attempts from each “deme” or interaction region, which we call *τ*_j_. In our tunable model, our time units are set such that the characteristic time between reproduction attempts is one. The rate of divisions that land within the interaction region is *p*, which sets the time scale of the logistic growth. Therefore, the characteristic saturation time of local regions is *α* = *p*, and as a zeroth-order estimate we have *τ*_s_ ≈ 1*/p*. This form is only useful for *p* close to one, because it ignores the effect of secondary events which land in the interaction region. As a result, the true dependence of *α* on *p* is weaker: *α* grows from 0.4 to 1 as *p* varies from zero to one (Fig. 12**a**). Therefore *τ*_s_ varies weakly from roughly 2.5 to one over the range of *p* values. For a more accurate estimate, we can use the phenomenological form *α* ≈ (1 + *p*)*/*2 ⇒ *τ*_s_ ≈ 2*/*(1 + *p*) which does not diverge as *p* → 0.

**FIG. 12.**
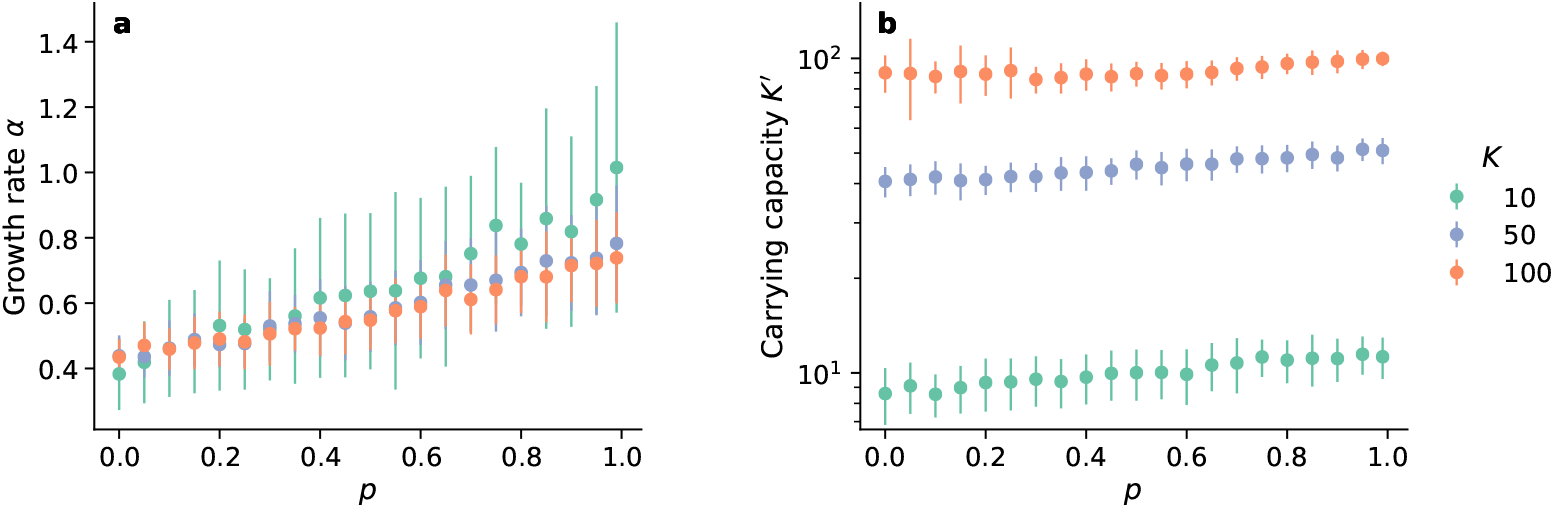
Fitted logistic growth parameters. **a.)** The growth rate increases with increasing short-range dispersal as expected. It does not depend on the carrying capacity because the growth rate is determined by the early growth of the population before the density regulation restricts population growth. **b.)** The fitted local carrying capacity increases slightly with increasing short-range dispersal. Regions often don’t saturate all the way to *K* as discussed above and shown in Fig. 10. We fit the logistic growth function to the saturation data of about 60 interaction regions across multiple simulations at each set of parameters. The points are averages and the error bars are standard deviations of the individual fits. This data comes from expansions with *μ* = 1.5 and is what formed the saturation times reported in Fig. 4c.

When regions have reached local saturation, the rate at which each interaction region sends out long-range jumps is *K*(1 − *p*). If we assume that the expansion is driven by jumps out of regions that have reached saturation, we have *τ*_j_ = 1*/*(*K*(1 − *p*)). Therefore, the condition *τ*_j_ ≫ *τ*_s_ reduces to

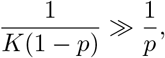

or

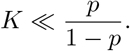

According to this criterion, most of our simulations explicitly do not satisfy the separation of time scales assumed in the lattice model.

### C. Logistic growth description of population dynamics within interaction regions

We started the logistic growth measurements by searching for sufficiently isolated individuals. To find individuals worthy of tracking, we searched at the end of every generation for individuals who had no one else within a distance of 10*r*_b_. Those individuals must have dispersed a long distance. We searched at the end of every generation until we found at least a predetermined minimum number of isolated individuals at the same time. We required several at the same time purely for convenience on the data processing side; these measurements could just as well have been gathered one at a time as we found the isolated individuals. Nevertheless, once we found the isolated individuals, we recorded the number within their interaction regions every generation until the end of the simulation. The local saturation data was then used to fit for logistic growth parameters. We fit to the logistic function of the form

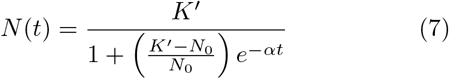

where *N* (*t*) is the population at time *t, K*′ is the local carrying capacity, *α* is the growth rate, and the initial population is *N*_0_ = 1. We used SciPy’s curve fit() function to make the fits and obtain *K*′ and *α*. We performed the fits on all individual interaction regions around the initially isolated individuals that we found that filled up to at least 60% of the local carrying capacity *K*. Average values and standard deviations are shown in SI Fig. 12.

We computed the saturation time for an interaction region by setting the population size in Eq. (7) equal to *K*′ − 1 and then solving for *t*, which leads to

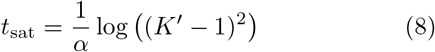

In addition to the dominant dependence ∼ 1*/α*, where *α* is itself proportional to *p* (see SI Section VI B), we find a logarithmic dependence of the saturation time on the local carrying capacity, which arises from the discrete nature of the local population within a deme. We computed the saturation time for every individual interaction region for which we fit the logistic growth function, using values of *α* and *K*′ from the fits to the logistic function. We report averages and standard deviations at *μ* = 1.5 in Fig. 4c.

For the local heterozygosity measurements, every individual in the initial population had a unique allele. We tracked the heterozygosity in the interaction regions of the same individuals for whom we measured logistic growth as described above (i.e. initially isolated individuals). The heterozygosities reported in Fig. 5 are averages of heterozygosities measured across typically about 50 separate interaction regions in the final generation of simulations and gathered from initially isolated individuals in multiple different simulations.

### D. Quantitative assessment of time-doubling hierarchy

We assessed the validity of simulation run times using the consistency condition ℓ(*t*)^*d*+*μ*^ ∼ *t*ℓ(*t/*2)^2*d*^ (eq. 1). The consistency condition is only valid after enough time has elapsed for long dispersal distances to be the driving factor behind a colony’s growth [16]. It is necessary to avoid the early times when applying the consistency condition, such as when estimating the kernel exponent as in Fig. 8. Colony growth remains self-consistent once the consistency condition becomes valid. For simulations that ran for *T* time steps, the values of *t* we used when applying the consistency condition ran from *T/*2 to *T*, so the values of *t/*2 ran from *T/*4 to *T/*2. The first data point we used with the consistency condition is marked with a red × in SI Fig. 13. We need at least a handful of data points after the first one to check for agreement with the consistency condition and to estimate the kernel exponent. Our run times were just enough at the lowest probabilities of local dispersal and gave us many useful data points at high local dispersal. We compared our data at *μ* = 2 against Eq. (1) in Fig. 7; analogous plots at *μ* = 1.5 and 2.5 are shown in SI Fig. 14.

**FIG. 13.**
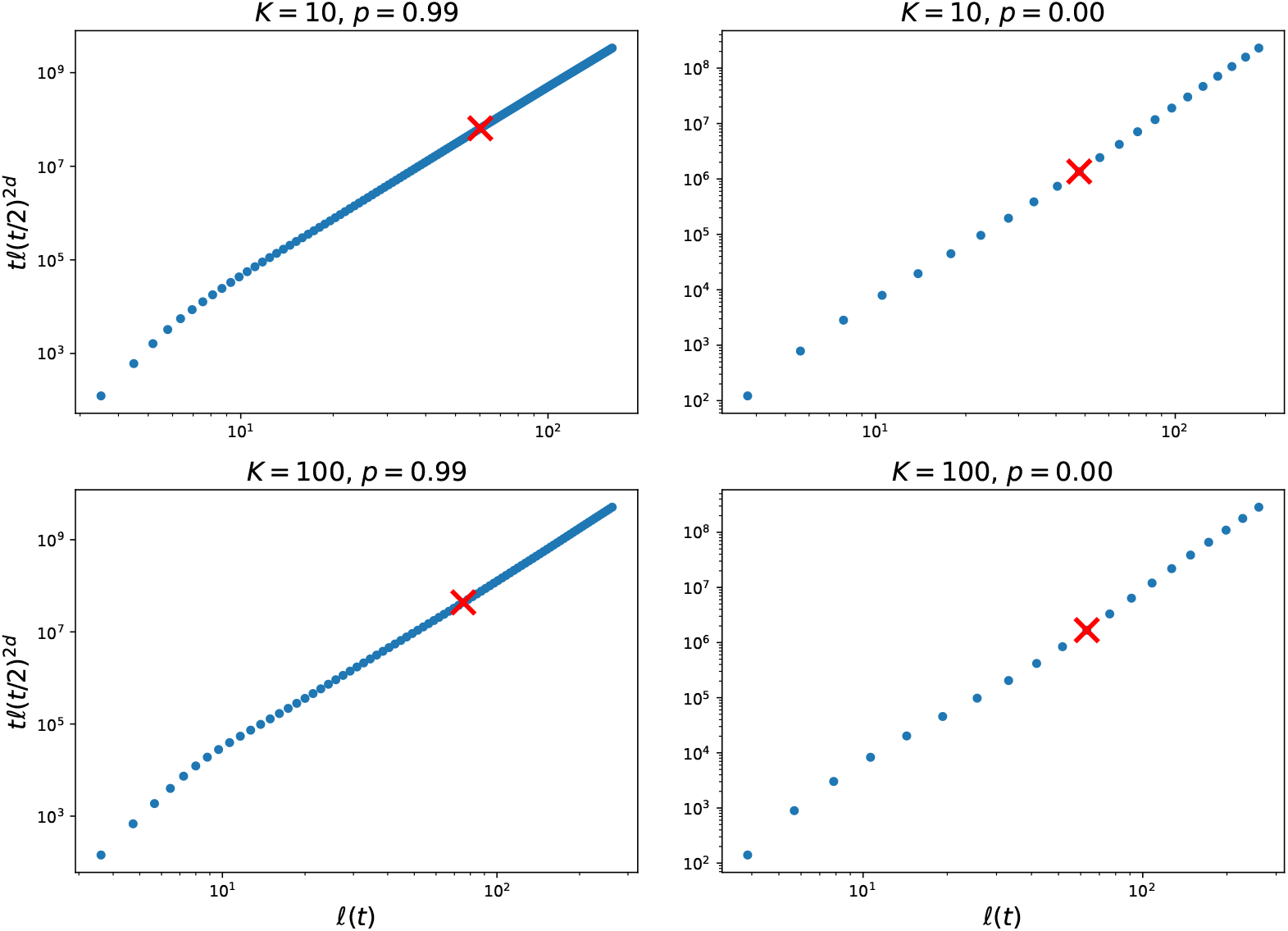
We show *t*ℓ(*t/*2)^2*d*^ plotted against ℓ(*t*) from average growth curves at all sets of parameters that went into figure 7. The dots are points from all even-numbered time steps. Simulations with high values of *p* require many more time steps to reach a given population size than those with low values of *p*. We use only the points after the red × in figure 7 and for inferring the kernel exponent as in figure 8. For simulations to be “long enough,” we needed at least a handful of points once the growth became self-consistent (i.e. linear on these plots). Using average growth curves from expansions at intermediate probabilities of local dispersal result in plots somewhere between these two extremes: more data points in the linear sections than the *p* = 0 case but not as many as in the *p* = 0.99 case.

The expansions at high local dispersal require many more time steps to reach the predetermined population threshold necessary to end simulations than those at low local dispersal. Expansions at high local dispersal take longer and grow slower than those at low local dispersal since offspring are much more likely to land near their parent in regions that may already be saturated, which means the times *t* and sizes ℓ(*t/*2) used here are higher at high local dispersal. A scaling factor of *A* ≈ 20 was needed to raise the points at low local dispersal to the level of those at high local dispersal. Bringing them together highlights the difference in power laws (slopes) between the sets of points at each value of *K*.

We found the inferred kernel exponent *μ*_i_ from the growth curves by fitting *B*ℓ(*t*)^*ν*^ to the quantity *t*ℓ(*t/*2)^2*d*^ using SciPy’s curve fit() function. We obtained values for both the prefactor *B* and the exponent *ν*, but only the exponent was of any interest for estimating the kernel exponent from the growth curves. We estimated the kernel exponent by performing this fit using data from only a later subset of the time steps as discussed in the previous paragraph. We then compute the inferred exponent as *μ*_i_ ≡ *ν* − *d*. We computed *μ*_i_ using all available growth curves (typically about 200 at any given set of parameters) to get the averages and confidence intervals reported in Fig. 8.

The exact value of *μ*_i_ somewhat depends on the fit method. For comparison, we repeated the process of extracting *μ*_i_ by finding the best fit line to the relevant data in log-log space, where the exponent could be found from the inferred slope. These two values would exactly match if we had infinitely long simulations that had perfectly converged to constant power laws, but in practice that is not the case. There is often a slight difference between the average values of *μ*_i_ from the two procedures, as shown at the example parameters in SI Fig. 15. However, we take the generally overlapping error bars as a signal that it’s safe to proceed with our inferred values. This sort of comparison could be used as a test of whether or not population growth has converged to the expected time-doubling hierarchy: consistent gaps between error bars are a warning that simulations may not be long enough. This test led us to run longer simulations to generate the data shown at *p* ≤ 0.5 in SI Fig. 15 and the corresponding data points in Fig. 8. The longer simulations ran until they reached population sizes of 30 million individuals, triple the size of our usual cutoff for simulations with *K* = 100.

**FIG. 14.**
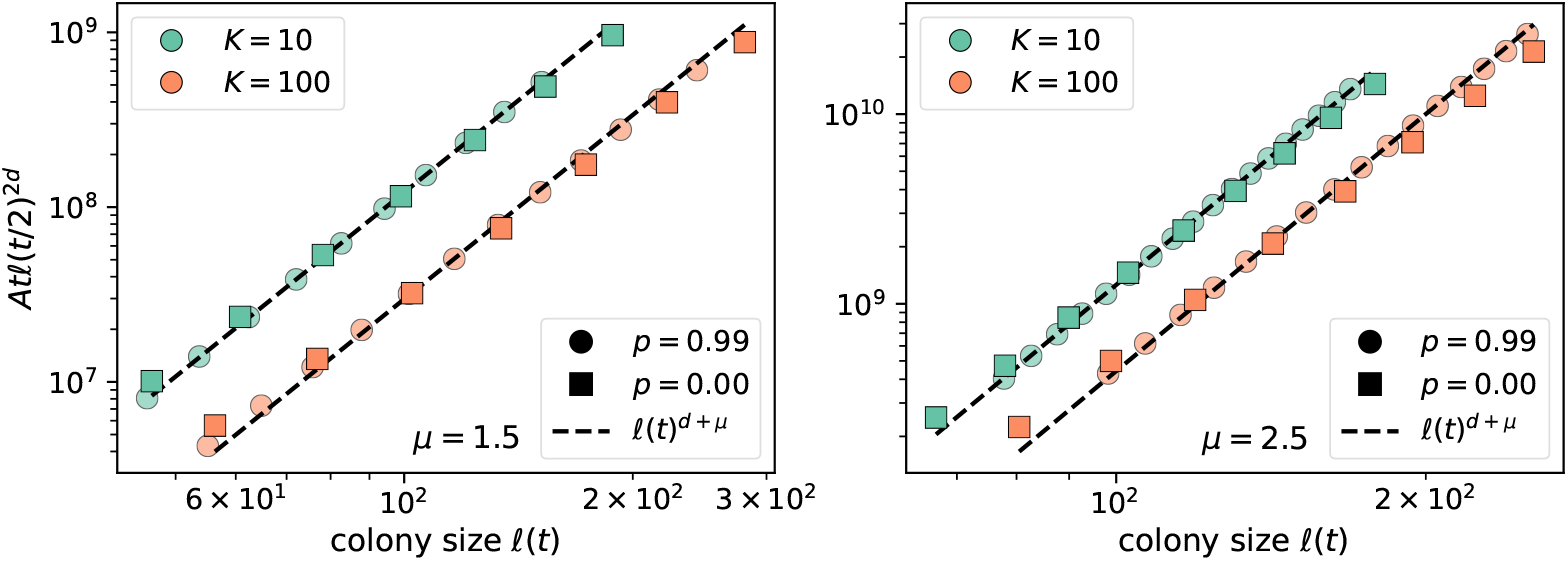
Analogous to Fig. 7.

**FIG. 15.**
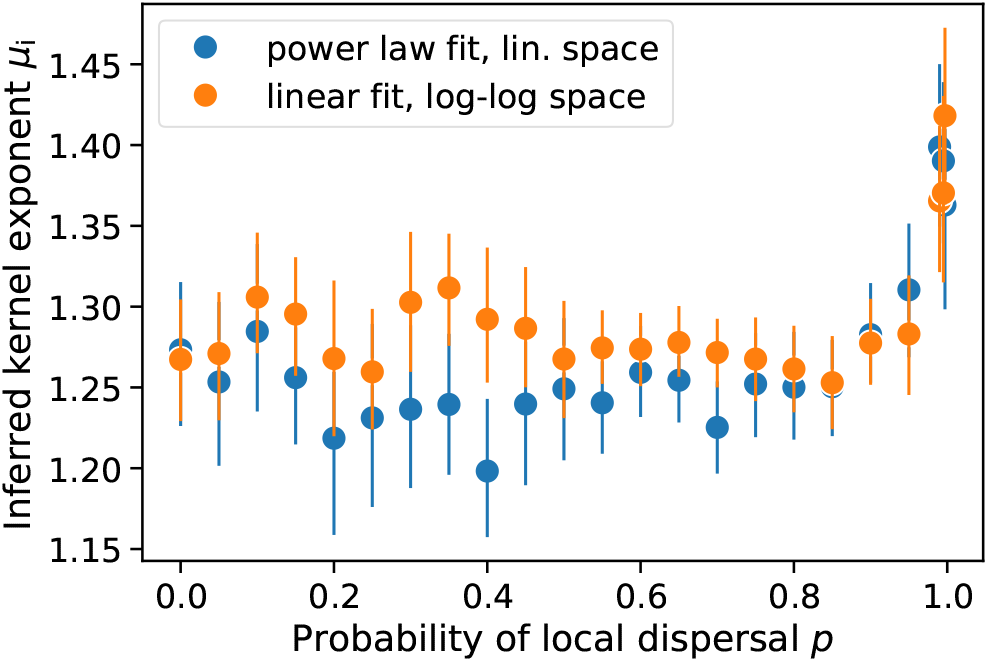
Example comparison between inferring kernel exponents by directly fitting power laws in linear space and fitting for the slope in log-log space. This example data comes from growth curves with *K* = 100 and *μ* = 1.5 (same data as the orange points in the left panel of Fig. 8); analogous plots look similar at other pairs of parameters. Each point represents the mean of the individual inferences from roughly 50 independent simulations below *p* = 0.5 and roughly 150 simulations above *p* = 0.5. The error bars are the 95% confidence interval of the distribution of bootstrapped mean inferred kernel exponents.

**FIG. 16.**
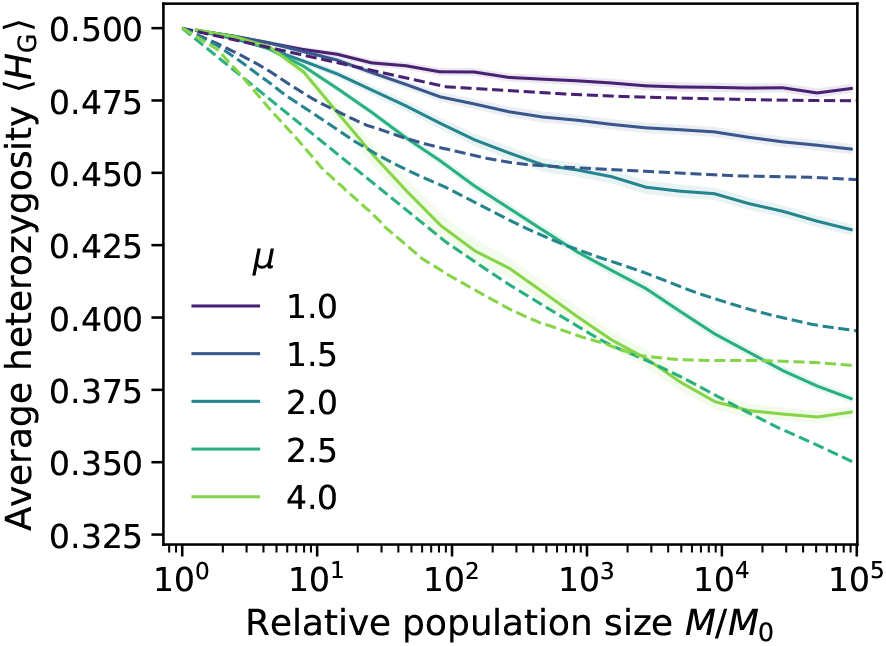
Direct comparison of heterozygosity evolution in the continuum simulations with fast local dynamics (*K* = 10, *p* = 0.5; solid lines and shading are same as in Fig. 9**a**) to that of the lattice-based simulations reported in Ref. 27 (dashed lines). In both cases, the initial population had a 50/50 mix of two alleles (initial heterozygosity of 0.5). Kernel exponents match at all values except *μ* = 2.5 (continuum), for which the same color refers to *μ* = 2.4 (lattice).

### E. Reporting the evolution of global heterozygosity

There are multiple reasonable ways to compute and display the global heterozygosity as a function of the growing population size as in Fig. 9 and SI Fig. 16; here we discuss some options and justify our choice. The true independent variable in our simulations is time. Every time step consists of offspring generation and dispersal followed by density regulation as discussed in Section III and SI Section VI A. Population size and heterozygosity are recorded at the end of each time step, after individuals have been removed from the population if their birthplaces are too densely occupied. This suggests that the “ground truth” for reporting the evolution of global heterozygosity might be plots of ⟨*H*_G_⟩ versus time, where averages and standard errors are computed with all available data at a given time step.

However, for generalizing results or comparing with the results of Ref. 27, it would be useful to compute ⟨*H*_G_⟩ as a function of the population size. One method of doing this would be to compute the averages ⟨*H*_G_⟩ and ⟨*M/M*_0_⟩ each time step. This method ignores what can be significant variation in population growth rates between individual simulations and generates points whose horizontal and vertical coordinates in the plots of Fig. 9 are both functions of time.

We sought to compute ⟨*H*_G_⟩ directly as a function of population size by generating binned population sizes and computing the average heterozygosity from all available simulation time steps where the population size was within a given bin. This means that a single simulation can contribute to a given data point on the ⟨*H*_G_⟩ versus *M/M*_0_ curve multiple times or not at all depending on how many time steps the population size was within that bin in that simulation. We used the `R` function `cut()` to place population sizes within 20 bins of equivalent width in logarithmic space, thus generating equally spaced data points for Fig. 9 and SI Fig. 16.

We use the standard error of the mean to estimate our uncertainty in ⟨*H*_G_⟩. A consequence of the binning procedure is that standard errors of the mean heterozygosity get vanishingly small in Fig. 9 and SI Fig. 16, despite the fact that the heterozygosity trajectory can vary quite a bit between individual simulations. The bins in those figures often consist of multiple data points from each simulation, especially for the bins at larger population sizes. Even though we generally have 200-400 independent simulations at each parameter combination shown in those figures, the points in the figures are often averages of thousands of data points that fall within each bin, resulting in nearly invisible uncer<u>ta</u>inty bands since the standard error of the mean is 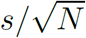 where *s* is the sample standard deviation and *N* is the number of samples. Such a large number of samples within each bin gives us a small uncertainty in our estimate of the average ⟨*H*_G_⟩.

### F. Direct comparison to lattice model

We used data for lattice-based simulations from Ref. 27 to compare results between continuum simulations at parameter values *K* = 10, *p* = 0.5 where the averaged normal heterozygosity is less than one (see Fig. 5), approximating local founder-takes-all, and lattice-based simulations where founder-takes-all is imposed at the deme level. The initial conditions in the two types of simulations were not exactly matched: both began with a 50/50 mix of two alleles, but the continuum simulations began with typically about 80 individuals near the origin (SI Section VI A) while the lattice-based simulations began with 111 occupied demes packed in a disc around the origin. Note that a deme is roughly a discrete analogue of an interaction region, so the continuum simulations’ 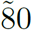 individuals correspond to roughly 80*/K* = 8 occupied demes. Another discrepancy is that Ref. 27 did not generate data at *μ* = 2.5, so we include their data from *μ* = 2.4 as a comparison with our *μ* = 2.5 data.

We observe that the difference in initial conditions leads to different dynamics at early times/small population sizes. In the continuum simulations, most of the early dynamics involves local events which mix and even out the starting population near the origin, and significant changes in heterozygosity only kick in when the population has reached ten times its initial size. By contrast, the lattice simulations only included long-range jumps, and the heterozygosity begins to fall earlier. This discrepancy leads to early differences in the observed heterozygosities between the two sets of models. However, the later trends, especially the contrast between a quick saturation of heterozygosity to a constant value at *μ* = 1 as opposed to a persistent decay for *μ* = 2.5 and a decay followed by a delayed saturation for *μ* = 4.0, are successfully captured by the lattice model. The quantitative discrepancy between the lattice and continuum values of ⟨*H*_G_⟩ is largest at *μ* = *d* = 2, which is a special point for the underlying dynamics that leads to extremely slow changes in the heterozygosity [16, 27]; we hypothesize that the small discrepancy in the initial conditions persists the longest at this special kernel exponent.

We also observe that the continuum ⟨*H*_G_⟩ at large population sizes is higher than that from lattice-based simulations for all jump-driven kernels (*μ* < 3). This is consistent with the observation that while local heterozygosity is small in the continuum simulations, it is not zero for the chosen parameter values of *K* = 10, *p* = 0.5 (left panel in Fig. 5) and the slight deviations from local founder takes all promote higher heterozygosity compared to the strict founder-takes-all assumption of the lattice model.

## References

[1] P. A. Delcourt and H. R. Delcourt, Long-term forest dynamics of the temperate zone (Springer (New York, NY), 1987).

[2] C. D. Thomas and J. J. Lennon, Nature 399, 213 (1999).

[3] L. D. Zeidberg and B. H. Robison, Proc. Natl. Acad. Sci. U.S.A. 104, 12948 (2007).

[4] S. D. Ling, Oecologia 156, 883 (2008).

[5] B. L. Phillips, G. P. Brown, J. K. Webb, and R. Shine, Nature 439, 803 (2006).

[6] C. Robertson, T. A. Nelson, D. E. Jelinski, M. A. Wulder, and B. Boots, J. Biogeogr. 36, 1446 (2009).

[7] V. Sousa, S. Peischl, and L. Excoffier, Curr. Opin. Genet. Dev. 29, 22 (2014).

[8] G. R. Walther, E. Post, P. Convey, A. Menzel, C. Parmesan, T. J. Beebee, J. M. Fromentin, O. Hoegh-Guldberg, and F. Bairlein, “Ecological responses to recent climate change,” (2002).

[9] L. Excoffier, M. Foll, and R. J. Petit, Annual Review of Ecology, Evolution, and Systematics 40, 481 (2009).

[10] C. A. Edmonds, A. S. Lillie, and L. L. Cavalli-Sforza, Proc. Natl. Acad. Sci. U.S.A. 101, 975 (2004).

[11] S. Klopfstein, M. Currat, and L. Excoffier, Mol. Biol. Evol. 23, 482 (2006).

[12] O. Hallatschek, P. Hersen, S. Ramanathan, and D. R. Nelson, Proc. Natl. Acad. Sci. U.S.A. 104, 19926 (2007), arXiv:0812.2345.

[13] O. Hallatschek and D. R. Nelson, Theoretical population biology 73, 158 (2008).

[14] L. Excoffier and N. Ray, Trends in Ecology & Evolution 23, 347 (2008).

[15] O. Hallatschek and D. R. Nelson, Evolution 64, 193 (2010), arXiv:0810.0053.

[16] O. Hallatschek and D. S. Fisher, Proceedings of the National Academy of Sciences of the United States of America 111, E4911 (2014), arXiv:1403.4639.

[17] M. Baguette, T. G. Benton, and J. M. Bullock, Dispersal ecology and evolution (Oxford University Press, 2012).

[18] R. Nathan, Science 313, 786 (2006).

[19] S. Hotaling, D. H. Shain, S. A. Lang, R. K. Bagley, L. M. Tronstad, D. W. Weisrock, and J. L. Kelley, Proceedings of the Royal Society B 286 (2019), 10.1098/RSPB.2019.0983.

[20] M. Chinazzi, J. T. Davis, M. Ajelli, C. Gioannini, M. Litvinova, S. Merler, A. Pastore y Piontti, K. Mu, ssi, K. Sun, C. Viboud, X. Xiong, H. Yu, M. Elizabeth Halloran, I. M. Longini, and A. Vespignani, Science 368, 395 (2020).

[21] R. A. Nichols and G. M. Hewitt, Heredity 72, 312 (1994).

[22] K. M. Ibrahim, R. A. Nichols, and G. M. Hewitt, Heredity 77, 282 (1996).

[23] V. Le Corre, N. Machon, R. J. Petit, and A. Kremer, Genet. Res. 69, 117 (1997).

[24] M. A. Lewis and S. Pacala, Journal of mathematical biology 41, 387 (2000).

[25] L. U. Wingen, J. K. Brown, and M. W. Shaw, Genetics 177, 435 (2007).

[26] R. Bialozyt, B. Ziegenhagen, and R. J. Petit, Journal of Evolutionary Biology 19, 12 (2006).

[27] J. Paulose and O. Hallatschek, Proceedings of the National Academy of Sciences, 201919485 (2020).

[28] M. Kot, M. A. Lewis, and P. Van Den Driessche, Ecology 77, 2027 (1996).

[29] J. M. Bullock, L. Mallada González, R. Tamme, L. Götzenberger, S. M. White, M. Pärtel, and D. A. Hooftman, Journal of Ecology 105, 6 (2017).

[30] K. S. Korolev, M. Avlund, O. Hallatschek, and D. R. Nelson, Reviews of Modern Physics 82, 1691 (2010), arXiv:0904.4625v2.

[31] J. Felsenstein, The American Naturalist 109, 359 (1975).

[32] N. H. Barton, J. Kelleher, and A. M. Etheridge, Evolution 64, 2701 (2010).

[33] N. H. Barton, A. M. Etheridge, and A. Véber, Journal of Statistical Mechanics: Theory and Experiment 2013, P01002 (2013).

[34] B. K. Epperson, Geographical genetics (MPB-38) (Princeton University Press, 2003).

[35] C. J. Battey, P. L. Ralph, and A. D. Kern, Genetics 215, 193 (2020).

[36] N. Ray and L. Excoffier, Molecular Ecology Resources 10, 902 (2010).

[37] C. E. Amorim, T. Hofer, N. Ray, M. Foll, A. Ruiz-Linares, and L. Excoffier, Heredity 118, 135 (2017).

[38] P. Ralph and G. Coop, Genetics 186, 647 (2010).

[39] J. Paulose, J. Hermisson, and O. Hallatschek, PLoS Genetics 15 (2019), 10.1371/journal.pgen.1007936.

[40] B. C. Haller and P. W. Messer, Molecular Biology and Evolution 36, 632 (2019).

[41] J. Kelleher, K. R. Thornton, J. Ashander, and P. L. Ralph, PLOS Computational Biology 14, 1 (2018).

[42] N. Berestycki, A. M. Etheridge, and A. Véber, Annales de l’Institut Henri Poincaré, Probabilités et Statistiques 49, 374 (2013).

[43] T. Smith and D. B. Weissman, bioRxiv, doi:10.1101/2020.06.24.168211 (2020).

[44] A. J. Nicholson, Australian journal of Zoology 2, 9 (1954).

